# A PROPPIN links V-ATPase assembly to endocytic membrane dynamics in malaria parasites

**DOI:** 10.64898/2026.04.14.718606

**Authors:** Yasmin Schmitz, Moumita Sengupta, Carola Schneider, Tanja Ziesmann, Franziska Hellmold, Ute Distler, Rudolph Reimer, Joachim Michael Matz

## Abstract

Malaria parasites replicate inside red blood cells, degrading hemoglobin within a specialized digestive vacuole. Efficient hemoglobin processing is essential for parasite survival and influences antimalarial drug susceptibility. The vacuole constantly fuses with incoming hemoglobin-filled vesicles, yet the mechanisms that balance cargo influx with membrane homeostasis remain unclear. Here, using conditional reverse genetics, quantitative live-cell imaging, and 3D electron microscopy, we characterize the autophagy-related protein 18 of *Plasmodium falciparum* (*Pf*ATG18) as a key regulator of vacuolar membrane dynamics. Loss of *Pf*ATG18 caused vacuole fragmentation, accumulation of hemoglobin-filled vesicles, and parasite death. These defects were preceded by broad architectural destabilization of the parasite’s V-ATPase, a proton pump controlling organelle acidification and the vacuole’s fusion–fission equilibrium. Direct interference with its membrane sector phenocopied *Pf*ATG18 deficiency. We found that *Pf*ATG18 does not interact directly with the proton pump but instead associates with a putative V-ATPase assembly factor and with complexes regulating phosphoinositide balance and vesicle trafficking. The breadth of these interactions indicates a multifaceted role at the vacuolar membrane and a regulatory influence on V-ATPase mediated through associated protein machinery. Although a point mutation in *Pf*ATG18 has been linked to artemisinin resistance, its complete knockout did not decrease sensitivity, but rather hypersensitized ring-stage parasites to dihydroartemisinin. Together, these findings establish *Pf*ATG18 as a central regulator of endocytic membrane homeostasis, essential for V-ATPase function and asexual parasite proliferation in the human blood.

## INTRODUCTION

Parasites of the genus *Plasmodium* are the causative agents of malaria. Within the human host, they replicate in erythrocytes, residing within a parasitophorous vacuole membrane (*1*). In this intracellular niche, the parasites progress through the ring, trophozoite and schizont stages before lysing the surrounding membranes to release their progeny. During intraerythrocytic development, the parasite catabolizes up to 80% of the erythrocyte cytoplasm, which is internalized at the cytostomes, invaginations of the parasite plasma membrane and the parasitophorous vacuole membrane (*2–4*). Cytostomes generate double-membrane-bound hemoglobin transport compartments (HTCs) that deliver host cell cytosol to an acidic, lysosome-like digestive vacuole (DV) (*4, 5*). Within the DV, hemoglobin is proteolytically degraded into small peptides and free heme (*5*). The peptides are exported into the parasite cytosol for further processing (*6, 7*), while heme is sequestered into bioinert hemozoin crystals, protecting the cell from oxidative damage (*8, 9*). Hemoglobin processing and heme biomineralization are essential for parasite survival in the blood, and many antimalarial drugs depend on hemoglobin catabolism for their activity. These include 4-aminoquinolines such as chloroquine, which interfere with hemozoin formation (*10, 11*), and artemisinin-based compounds, which are activated by heme (*12–14*). Consequently, the kinetics of heme release strongly influence drug efficacy, and mutations that impair endocytosis or proteolysis can confer resistance (*15–18*).

During each intraerythrocytic cycle, the DV is formed *de novo* from small acidified vesicles (*19*). In the most virulent malaria parasite species, *Plasmodium falciparum*, these vesicles coalesce into a single large organelle that undergoes frequent membrane fusion events to accommodate the constant influx of endocytic cargo (*5, 19*). Fusion between the outer HTC membrane and the digestive vacuole membrane (DVM) delivers a single-membrane-bound hemoglobin package into the DV matrix, where it is lysed to release the cargo (*2, 20, 21*). Similar to other eukaryotic systems, endocytic membrane fusion in malaria parasites depends on a variety of trafficking factors, including small GTPases (*22*), membrane tethering complexes (*23*), and likely SNARE proteins (*2*). The endocytic signature lipid phosphatidylinositol 3-phosphate (PI3P) plays a critical role in this process by recruiting trafficking factors to the membranes of both HTCs and the DV (*24–27*). In *Plasmodium*, PI3P is synthesized by the class III phosphoinositide 3-kinase (PI3K) Vps34, likely in association with regulatory subunit Vps15, and its genetic or pharmacologic disruption has been shown to impair HTC-DVM fusion (*28, 29*).

The continuous delivery of HTC membranes to the DV necessitates mechanisms that prevent excessive expansion of the vacuole. In other systems, this is achieved by transferring lipids to a receiving organelle at membrane contact sites (*30*) or by shaping the membrane into tubules or vesicles that are then pinched off with the help of scission proteins (*31*). The latter was shown to depend on phosphatidylinositol 3,5-bisphosphate [PI(3,5)P_2_], which is produced from PI3P by the Fab1/PIKfyve complex (*32*). This assembly consists of the kinase Fab1/PIKfyve, the scaffold protein Vac14, and the phosphatase Fig4, which can reverse the reaction (*33*). The balance between PI3P and PI(3,5)P_2_ therefore plays a critical role in regulating delivery and recycling of membrane material along the endocytic pathway. Although *Plasmodium* parasites encode all core subunits of the Fab1/PIKfyve complex, their functions remain uncharacterized, and the production of PI(3,5)P_2_ by the parasite has yet to be demonstrated (*34*).

The molecular mechanisms that govern membrane equilibrium at the *Plasmodium* DV are still poorly understood; however, we recently found that the V-ATPase plays a key role in this process (*20*). The V-ATPase is a hetero-multimeric proton pump that acidifies the DV matrix. It consists of an ATP-hydrolyzing V_1_ sector facing the cytosol and a membrane-embedded V_0_ sector responsible for proton translocation (*35*). The two sectors are reversibly coupled by a regulatory scaffold, called the RAVE complex in yeast or Rabconnectin-3 (Rbcn3) in mammals (*36*). Beyond its canonical function in vacuolar acidification, we found that the V_0_ sector is critical for maintaining DV integrity. Loss of either the V_0_a or V_0_c subunits caused dissociation of the V_1_ sector from the vacuolar membrane and triggered DV fission (*20*). Interestingly, genetic inactivation of the catalytic V_1_B subunit, which also led to V_1_ sector dissociation, did not compromise DV integrity, indicating that V-ATPase regulation of DVM dynamics is independent of the V_1_ sector and the catalytic activity of the complex (*20*).

While the V-ATPase appears to limit DV fission, other proteins may promote it in order to maintain vacuolar membrane homeostasis. Indeed, the *P. falciparum* DV can form autonomous membrane lobes that detach from the parent organelle (*19, 37*). A likely contributor to this process is the parasite’s autophagy-related protein 18 (*Pf*ATG18), a member of the β-propeller that bind polyphosphoinositides (PROPPIN) family, which has been implicated in endocytic membrane scission in other organisms. In *Saccharomyces cerevisiae*, ATG18 consists of WD40 repeats forming a seven-bladed β-propeller, with flexible loops extending from the blades (*38*) (Figure 1A). It is required for vacuole fission and exhibits membrane scission activity *in vitro,* which depends on insertion of its 6CD loop into membranes and on its binding to PI3P and PI(3,5)P_2_ (*39*). This activity is enhanced by interaction with the retromer complex, a protein coat that forms endosomal transport carriers (*40*). Disrupting retromer engagement of the mammalian ATG18 homolog, WD repeat domain phosphoinositide-interacting protein 1 (WIPI1), leads to accumulation of elongated endosomes, further supporting a role for the PROPPIN family in endocytic membrane fission (*40*). Importantly, this function of ATG18 is independent of its role in autophagy, which involves the recruitment of ATG2, the scramblase ATG9, and other autophagy machinery to facilitate lipid transfer from the endoplasmic reticulum (ER) to the phagophore (*40–42*).

**Figure 1.**
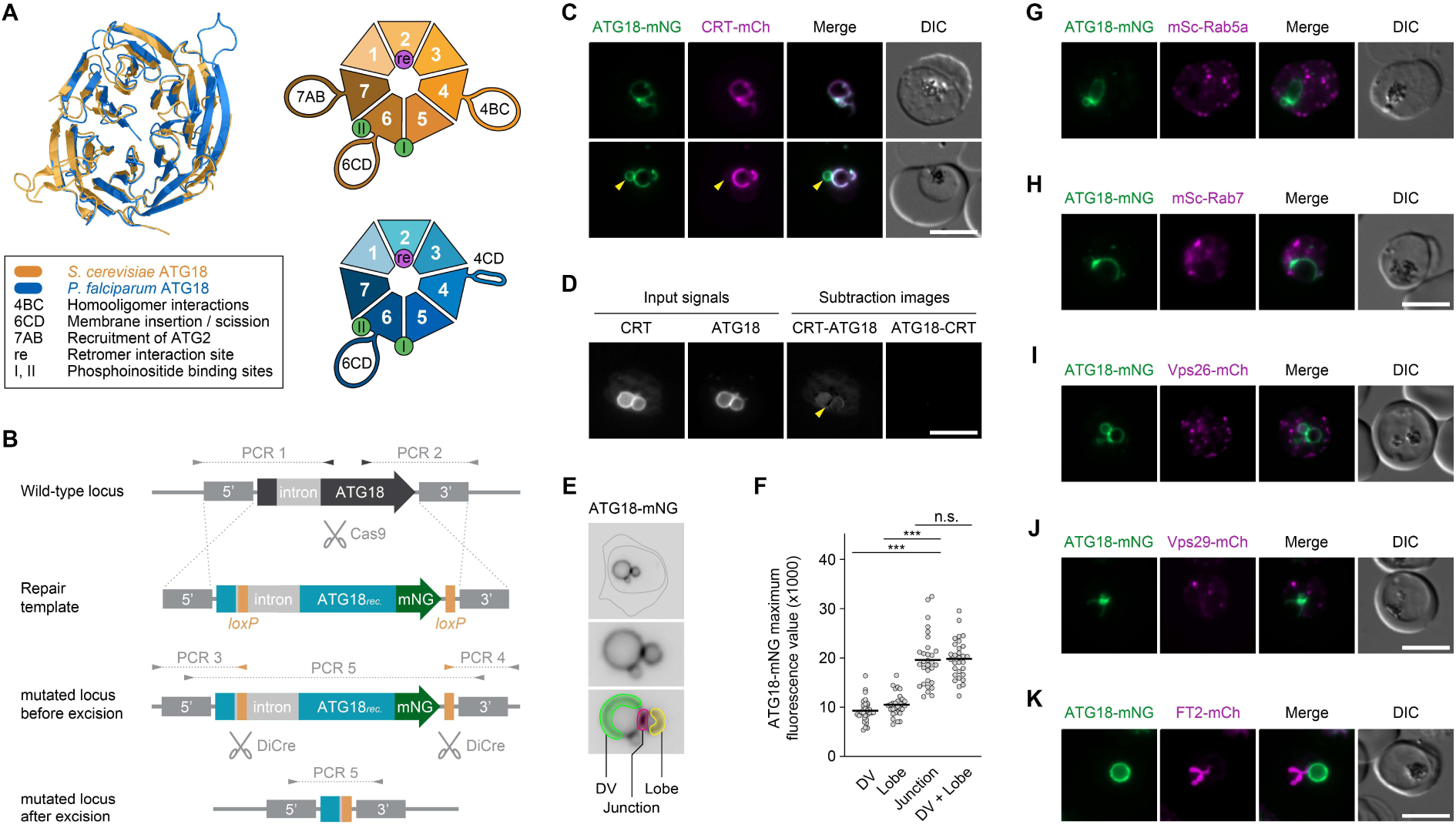
*Pf*ATG18 marks the DVM without junctional enrichment. **(A)** Structural organization of *Pf*ATG18. Alignment of the *Sc*ATG18 crystal structure (PDB: 6KYB; (*38*)) with the *Pf*ATG18 AlphaFold 3 model reveals a conserved seven-bladed β-propeller core. Loops were omitted due to absence in the crystal structure or low-confidence predictions. RMSD = 1.18 Å. Key structural features are highlighted in the schematics and legend. **(B)** Genetic strategy for tagging and conditional deletion of *Pf*ATG18. Cas9-mediated cleavage and homologous recombination yield *atg18-mNG:loxP* parasites expressing *loxP*-flanked recodonized (rec.) *Pf*ATG18 fused mNG. RAP-inducible DiCre excises the *loxP*-flanked sequence. Arrowheads and dotted lines indicate primers and expected PCR products. A second independent line (*atg18:loxP*) lacks the mNG tag. **(C)** *Pf*ATG18 colocalizes with CRT at the DVM and is occasionally observed at DV-adjacent compartments (arrowhead). **(D-F)** *Pf*ATG18 is not enriched at DV-lobe junctions. (D) Normalized CRT-mCh and *Pf*ATG18-mNG intensities were subtracted to reveal signals exclusive to each protein. The arrowhead marks the membrane junction. Maximum ATG18-mNG intensities at DV, lobe, and junction (E) showed no junctional accumulation beyond the sum of both membranes (F). Individual and mean values; one-way ANOVA with Tukey’s test; n = 30 parasites, 3 experiments. n.s., P > 0.05; ***, P < 0.001. **(G-K)** *Pf*ATG18 shows no overlap with fluorescently tagged early (G) or late (H) endosomal markers, retromer components (I,J) or an apicoplast transporter (K). Scale bars: 5 µm.

Structural homology modelling using AlphaFold3 suggests that the *P. falciparum* homolog of ATG18 retains the core architecture of other PROPPINs, including the seven-bladed β-propeller (Figure 1A). *Pf*ATG18 also features the 6CD loop required for membrane insertion and scission but lacks the 7AB loop, which typically mediates the recruitment of ATG2 during autophagy (*38*). This is consistent with the apparent absence of homologs for ATG2, ATG9 and other core autophagy machinery in *Plasmodium*, which has cast doubt on the existence of a canonical autophagy pathway in these highly divergent eukaryotes (*43*).

The localization and functions of *Pf*ATG18 remain uncertain. The protein has been observed at the DVM, with pronounced accumulations at the interface between the DV and its lobes (*25, 37*), as well as in vesicular compartments and in proximity to the parasite’s non-photosynthetic plastid, the apicoplast (*44, 45*). Reverse genetic experiments in *P. falciparum* and the related apicomplexan parasite *Toxoplasma gondii* have suggested a role for ATG18 in apicoplast inheritance (*44*). Nevertheless, multiple lines of evidence support a function for *Pf*ATG18 at the DVM: (1) it binds efficiently to PI3P *in vitro* and relocates to the parasite cytoplasm when its phosphoinositide-binding sites are mutated or when PI3P production is pharmacologically inhibited (*25*), (2) it interacts with the DVM-resident transporter multidrug resistance protein 1 (MDR1) in co-immunoprecipitation experiments (*25*), (3) its expression restores vacuolar fission in ATG18-deficient yeast mutants (*37*), and (4) a T38I point mutation in its coding sequence is linked to slower parasite clearance in patients treated with artemisinin-based antimalarials (*45, 46*), whose activation depends on hemoglobin catabolism in the DV (*12–14, 17, 18*).

The uncertainty surrounding the involvement of *Pf*ATG18 in autophagy, apicoplast inheritance and vacuolar membrane dynamics prompted us to undertake an in-depth functional characterization of this PROPPIN family member. Using conditional reverse genetics combined with quantitative live cell imaging and 3D electron microscopy, we demonstrate that *Pf*ATG18 function is critical for DV integrity and endocytic flux. Moreover, we identify a mechanistic link between *Pf*ATG18 and the parasite’s V-ATPase, showing that *Pf*ATG18 facilitates proper assembly of the proton pump to maintain vacuolar membrane homeostasis.

## RESULTS

### *Pf*ATG18 localizes to the DVM

To study the functions of *Pf*ATG18 during blood stage development, we first generated conditional *Pf*ATG18 knockout parasites on the genetic background of the dimerizable Cre recombinase (DiCre)-expressing B11 strain of *P. falciparum* (3D7) (*47*). Using Cas9-mediated genome editing, two *loxP* sites were introduced into the *Pf*ATG18 locus, one within an intron near the start codon and one downstream of the stop codon (*48*) (Figure 1B). These modifications were designed to allow for DiCre-mediated excision of 89% of the *Pf*ATG18 coding sequence upon addition of rapamycin (RAP). Two independent conditional knockout lines were generated; in the *atg18-mNG:loxP* line, *Pf*ATG18 was tagged with mNeonGreen (mNG) at the C-terminus, while in the *atg18:loxP* line, the protein was left untagged. Site-specific integration of the modifying DNA constructs into the parasite genome was confirmed by diagnostic PCR (Supplemental Figure S1).

To gain insights into the expression and localization of *Pf*ATG18, we studied *atg18-mNG:loxP* parasites by live fluorescence microscopy, which revealed abundant expression of the protein throughout the intraerythrocytic cycle (Supplemental Figure S2A). In young ring stages, *Pf*ATG18-mNG localized to a slender elongated structure. In more mature stages, the protein was consistently detected in a circular pattern around the central hemozoin cluster, suggestive of the DVM (Supplemental Figure S2A,B). We further engineered the *atg18-mNG:loxP* parasites to express mCherry (mCh)-tagged plasmepsin II (PM2) or chloroquine resistance transporter (CRT), markers of the vacuolar matrix and membrane, respectively. Live microscopy showed that *Pf*ATG18-mNG surrounded the PM2-mCh-positive DV matrix (Supplemental Figure S2C), and overlapped extensively with CRT-mCh at the DVM (Figure 1C). Only in very few instances did we observe in direct contact with the DV an additional *Pf*ATG18-mNG positive compartment harboring no CRT-mCh (Figure 1C). However, the low frequency of these observations indicates that *Pf*ATG18 occupancy of this compartment, or the compartment itself, is transient.

### *Pf*ATG18 is not enriched at DVM junctions

It was previously proposed that *Pf*ATG18 is specifically recruited to the membrane junctions between the DV and its lobes (*37*). To test this, we recorded *atg18-mNG:loxP + crt-mCh* parasites harboring lobed vacuoles. By performing a pixel-by-pixel subtraction of the normalized *Pf*ATG18-mNG and CRT-mCh input signals, we visualized the pools unique to each of the proteins. CRT-mCh showed unique weak signals in the parasite cytoplasm and in the vacuolar lumen, most likely corresponding to ER-resident trafficking intermediates and porphyrin-mediated autofluorescence, respectively (Figure 1D). Except for the rare transient structures mentioned above, *Pf*ATG18-mNG occupied no areas in the cell that were not also occupied by CRT-mCh. Importantly, the contact sites between the DV and its lobes displayed no specific enrichment of *Pf*ATG18-mNG (Figure 1D).

We then measured the maximum *Pf*ATG18-mNG fluorescence intensity at three different zones of the DVM: at the mother vacuole, at the lobe and at their shared junction (Figure 1E). The signal at the junctional sites was approximately twice that of the individual sub-compartments, as expected for close apposition of two identical membranes (Figure 1F). Accordingly, the signal intensity at the membrane junction was equal to the sum of the fluorescence values of both sub-compartments. Together, these observations indicate no enrichment of *Pf*ATG18 at the DV-lobe interface beyond that predicted by close membrane apposition.

### *Pf*ATG18 does not colocalize with endosomal, retromer or apicoplast markers

Motivated by previous reports of yeast and mammalian ATG18 homologs functioning in endosomal membrane homeostasis and retromer engagement (*40*), we sought to determine whether a pool of *Pf*ATG18 localizes to compartments along the endosomal pathway or to sites of retromer assembly. To that end, we expressed mCh or mScarlet (mSc)-tagged reporter proteins on the genetic background of the *atg18-mNG:loxP* parasite line for live colocalization analysis. We detected no convincing overlap of *Pf*ATG18-mNG with Rab GTPases of early and late endosomes (Figure 1G,H) or with the retromer components Vps26 and Vps29 (Figure 1I,J), which all displayed punctate or patchy distributions throughout the parasite cytoplasm. In contrast to earlier reports (*44*), we detected no association of *Pf*ATG18-mNG with the apicoplast (Figure 1K). Combined, our comprehensive localization analysis suggests that *Pf*ATG18 localizes exclusively to the DVM with rare transient recruitment to a compartment closely associated with, or belonging to, the DV.

### *Pf*ATG18 is essential for asexual blood stage development

To study the consequences of *Pf*ATG18 inactivation, we induced DiCre-mediated genomic excision of the *Pf*ATG18 coding sequence by supplementing tightly synchronized *atg18-mNG:loxP* parasite cultures with 20 nM RAP from the ring stage onward. This resulted in efficient excision of the *loxP*-flanked genomic sequence by the end of the first intraerythrocytic cycle, as shown by diagnostic PCR (Figure 2A). Western blot analysis and live cell imaging indicated that *Pf*ATG18-mNG protein was strongly depleted upon RAP treatment, resulting in only ∼6% of its physiological expression levels by 36 hours post invasion (hpi) (Figure 2B-D). By contrast, expression of CRT-mCh remained completely unaffected by the absence of *Pf*ATG18 in the corresponding double mutant (Figure 2E).

**Figure 2.**
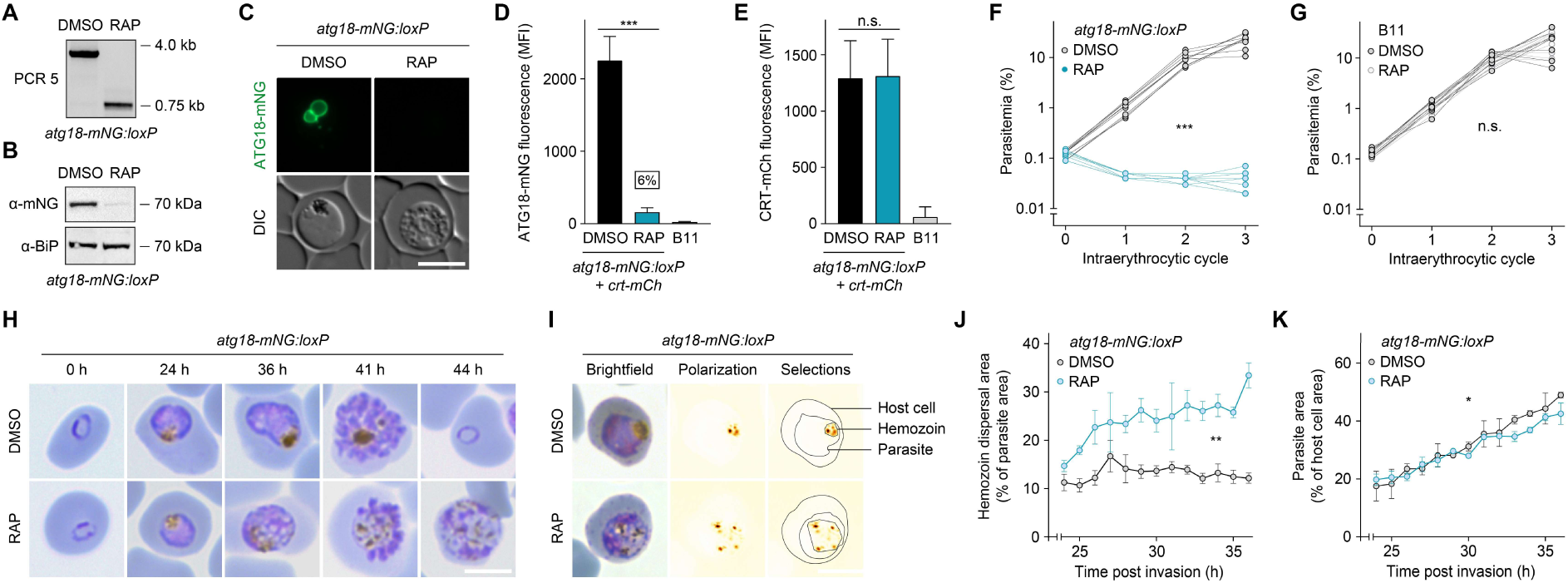
Loss of *Pf*ATG18 causes hemozoin dislocation and parasite death. Synchronized parasites were treated with DMSO or RAP from the ring stage onward. **(A)** RAP-induced excision of *loxP*-flanked genomic DNA in *atg18-mNG:loxP* parasites, confirmed by diagnostic PCR 5 (see Figure 1A). **(B-E)** RAP efficiently depletes *Pf*ATG18-mNG by 36 hpi, as shown by Western blot (B) and live-cell imaging (C). Quantitative fluorescence microscopy reveals strong reduction of *Pf*ATG18 (D), but not CRT (E), following RAP treatment. Parental B11 parasites serve as background control. Mean +/− SD; one-way ANOVA with Tukey’s test; n = 99 parasites, 3 experiments. n.s., P > 0.05; ***, P < 0.001. **(F-H)** *Pf*ATG18 is essential for asexual blood-stage development. RAP blocks replication of *atg18-mNG:loxP* parasites (F) but not B11 controls (G). Individual values; two-way ANOVA; n = 3 experiments with 3 technical replicates each. (H) Giemsa-stained *atg18-mNG:loxP* parasites throughout one intraerythrocytic cycle. **(I-K)** Loss of *Pf*ATG18 alters hemozoin distribution. (I) Giemsa-stained parasites imaged by brightfield and polarization microscopy. Hemozoin dispersal area (J) and parasite area (K) were quantified. Mean +/− SEM; two-way ANOVA; n = 99 parasites, 3 experiments. *, P < 0.05; **, P < 0.01. Scale bars: 5 µm.

In flow cytometry-assisted growth assays, dimethyl sulfoxide (DMSO)-treated *atg18-mNG:loxP* control parasites multiplied efficiently, whereas RAP treatment caused a complete halt in parasite replication (Figure 2F). Off-target effects of the inducing compound can be dismissed, as RAP supplementation did not measurably affect growth of the parental B11 strain (Figure 2G). Microscopic inspection of Giemsa-stained blood smears revealed that *Pf*ATG18-deficient parasites arrested their development at young schizont stage, which was accompanied by striking morphological abnormalities including cytoplasmic vacuolarization and a scattered distribution of hemozoin crystals throughout the parasite cell (Figure 2H). We thus conclude that *Pf*ATG18 is essential for asexual parasite replication in the blood.

### DVs lacking *Pf*ATG18 undergo fission but do not lyse

Using combined polarization and brightfield microscopy of Giemsa-stained blood smears (Figure 2I), we found that the area occupied by the hemozoin crystals remained relatively constant throughout development of DMSO-treated *atg18-mNG:loxP* control parasites between 24 and 36 hpi (Figure 2J). By comparison, hemozoin crystals in *Pf*ATG18-deficient parasites occupied a larger area already at trophozoite stage, and this area increased significantly over time. By 36 hpi, the hemozoin dispersal area of RAP-treated *atg18-mNG:loxP* parasites exceeded that of DMSO controls by 2.8-fold (Figure 2J). Notably, the drastic differences in subcellular hemozoin distribution did not correlate with a lack of parasite growth, as the area of *Pf*ATG18-deficient parasites increased steadily over time, albeit at a slightly reduced rate compared to DMSO controls (Figure 2K).

The scattered appearance of hemozoin crystals in in the absence of *Pf*ATG18 prompted us to study the morphology of the DV in DMSO and RAP-treated *atg18:loxP* parasites by live fluorescence microscopy of CRT-mNG. As expected, CRT localized to a single DV in DMSO control parasites. Upon inactivation of *Pf*ATG18, however, CRT was associated with a large number of vesicular compartments spread throughout the parasite, indicative of vacuolar fragmentation (Figure 3A). To clarify whether this coincided with a loss of membrane barrier function, we monitored PM2 localization, and found the signal to be restricted to the vacuolar fragments and absent from the parasite cytosol, indicating no breach of DVM integrity upon *Pf*ATG18 knockout (Figure 3B). Together, these findings suggest that, unlike its yeast and mammalian counterparts, *Pf*ATG18 does not promote vacuole fission, but rather acts to constrain it.

**Figure 3.**
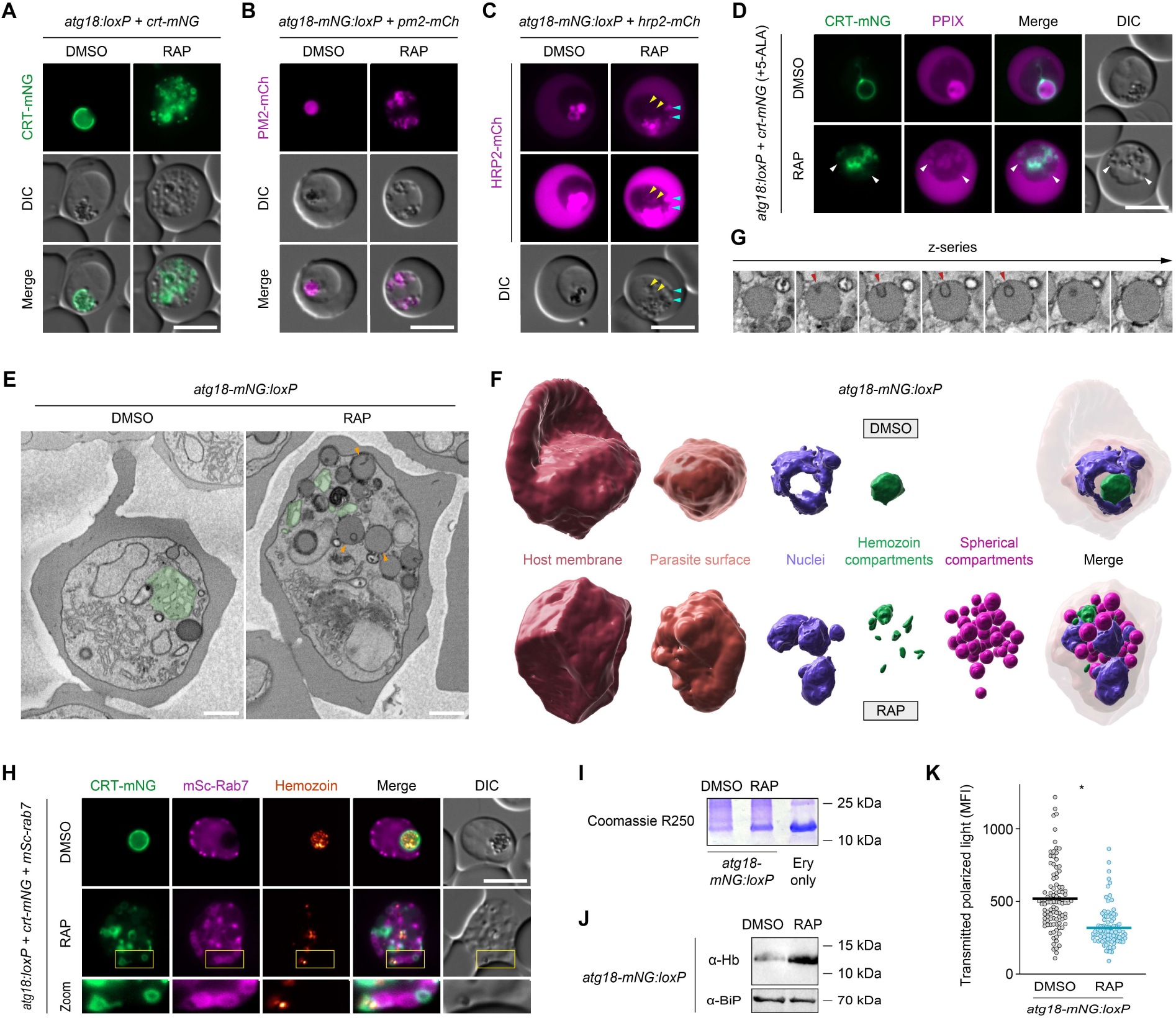
*Pf*ATG18 is required for DV maintenance and hemoglobin delivery. Synchronized parasites were treated with DMSO or RAP from the ring stage onward and analyzed at 36 hpi. **(A,B)** Loss of *Pf*ATG18 causes fission of the DVM without compromising its barrier function. Shown are fluorescently tagged CRT (A), and PM2 (B). **(C,D)** *Pf*ATG18-deficient parasites harbor two distinct populations of host cell cytosol-containing vesicles. (C) Localization of HRP2, a parasite protein exported into the host cell and subsequently reinternalized. Cyan arrowheads: bright compartments; yellow arrowheads: dim compartments. (D) *atg18:loxP + crt-mNG*-infected erythrocytes were treated with 5-aminolevulinic acid (5-ALA) to induce synthesis of fluorescent protoporphyrin IX (PPIX) in the host cell. White arrowheads: PPIX-positive compartments lacking CRT. **(E-G)** SBF-SEM reveals DV fragmentation and accumulation of spherical compartments in *Pf*ATG18-deficient parasites. Representative sections (E) and 3D reconstructions (F) highlight relevant organelles. (G) Z-series of a spherical compartment in a *Pf*ATG18-deficient parasite. Green: hemozoin-containing compartments; orange arrowheads: spherical compartments; red arrowheads: membrane invagination overlain by a second limiting membrane. Scale bars: 1 µm. **(H)** *Pf*ATG18-deficient parasites accumulate compartments positive for the late endosomal marker Rab7. **(I,J)** Loss of *Pf*ATG18 results in accumulation of undigested hemoglobin. Parasites released by saponin lysis were analyzed by SDS-PAGE, and hemoglobin (Hb) was visualized by Coomassie staining (I) or Western blotting (J). Ery: erythrocyte extract. **(K)** Loss of *Pf*ATG18 reduces hemozoin formation, as quantified by polarization microscopy. Individual and mean values; paired *t*-test; n = 99 parasites, 3 experiments. *, P < 0.05. Scale bars: 5 µm unless noted otherwise.

### Two distinct vesicle populations in *Pf*ATG18-deficient parasites

We then tested whether the vacuolar fragments of *Pf*ATG18-deficient parasites contained host cell cytosol by monitoring the localization of mCh-tagged histidine-rich protein II (HRP2), a soluble parasite protein that is exported into the host cell compartment and accumulates in the DV by endocytic uptake of erythrocyte cytosol (*49*). In DMSO control parasites, HRP2-mCh was found associated with the host cell compartment, but was most strongly concentrated in the DV matrix (Figure 3C). Upon inactivation of *Pf*ATG18, the vacuolar HRP2-mCh staining appeared fragmented and we observed several highly fluorescent vesicles that were clearly detached from the mother vacuole. We also detected a second population of vesicles in *Pf*ATG18-deficient parasites with much lower HRP2-mCh intensity, comparable to that of the host cell compartment (Figure 3C).

To determine whether these vesicles were also derived from the collapsed DV, we visualized internalized host cell cytosol in *atg18:loxP* parasites expressing CRT-mNG. In this experiment, we chose an alternative strategy for host cell cytosol labelling that exploits the erythrocyte’s capacity to synthesize the fluorescent heme precursor protoporphyrin IX (PPIX) upon supplementation with 5-aminolevulinic acid (*50*). As with HRP2-mCh, PPIX fluorescence was detected in the host cell cytosol and in the DV of DMSO control parasites (Figure 3D). Upon inactivation of *Pf*ATG18, the intraparasitic PPIX pool was distributed among numerous vesicular compartments. Not all of these structures were delineated by CRT-mNG, indicating that a considerable fraction of host cell cytosol-containing vesicles formed independently of vacuolar fission (Figure 3D).

### 3D ultrastructure of *Pf*ATG18-deficient parasites

The complexity of the *Pf*ATG18 knockout phenotype prompted us to study the ultrastructure of DMSO and RAP-treated *atg18-mNG:loxP* parasites by serial block-face scanning electron microscopy (SBF-SEM) (Supplemental Movies S1 and S2). Three-dimensional reconstruction confirmed that the DV of RAP-treated parasites fragments into numerous hemozoin-containing vesicles without signs of membrane lysis, with an average volume of 0.35 fL (Figure 3E,F; Supplemental Figures S3 and S4A; Supplemental Movies S3 and S4). In addition, these parasites exhibited a striking accumulation of spherical compartments with a mean volume of 1 fL (Supplemental Figure S4A). The electron backscatter signal of these compartments was slightly higher than that of the erythrocyte cytoplasm, suggesting they contain host cell hemoglobin (Supplemental Figure S4B). Consistent with this, previous studies using a membrane-impermeable fluorescent dextran have shown that host cell cytosol is concentrated within vesicular parasite compartments following endocytic uptake (*19*). In very few instances did we observe one or two of the spherical compartments also in DMSO-treated control parasites, indicating that they are physiological but transient. Although the resolution of SBF-SEM does not always allow clear discrimination of closely apposed membranes, we frequently observed inward folds of the spherical compartment envelope, occasionally overlain by a second membrane (Figure 3G).

The spherical compartments displayed a range of different membrane features, which we grouped into five categories: (1) slits protruding inward from the periphery, often from a funnel-like invagination; (2) slits terminating in an electron-dense vesicle; (3) peripherally associated electron-dense vesicles without a slit; (4) similarly sized sub-compartments divided by a membrane septum; and (5) electron-lucent inclusions (Supplemental Figure S5). Slits and electron-dense vesicles, either alone or in combination, were by far the most common features, while translucent inclusions were rare. All analyzed compartments harbored at least one of these membrane features, with some exhibiting multiple. By contrast, 3D-reconstructed cytostomal invaginations displayed no such modifications, suggesting that they are introduced during or after budding from the cytostome (Supplemental Figure S5).

Finally, we note that, despite previous suggestions of *Pf*ATG18 being involved in apicoplast maintenance, the organelle exhibited comparable size and morphology in both DMSO– and RAP-treated parasites (Supplemental Figure S3).

### Defective HTC fusion blocks hemoglobin processing in *Pf*ATG18-deficient parasites

We hypothesized that the spherical compartments in *Pf*ATG18-deficient parasites represent late HTCs unable to fuse with the DV, resulting in an inability to process hemoglobin. To test this, we first examined the localization of mSc-tagged Rab7, a late endosomal marker (Figure 3H). In DMSO control parasites, several punctate Rab7-positive compartments were observed. In contrast, *Pf*ATG18-deficient parasites harbored an additional population of enlarged Rab7-decorated compartments, often positioned near CRT-positive DV fragments. This pattern was not observed with the early endosomal marker Rab5A (Supplemental Figure S6), together indicating that the spherical compartments accumulating in the absence of *Pf*ATG18 are matured HTCs that fail to undergo fusion.

To confirm the accumulation of unprocessed HTC cargo, protein extracts from saponin-released parasites were analyzed by SDS-PAGE. Coomassie staining revealed that *Pf*ATG18-deficient parasites were enriched in a protein of ∼15 kDa, which comigrated with the most abundant protein species detected in pure erythrocyte extracts (Figure 3I). Western blot analysis confirmed that this protein was hemoglobin (Figure 3J). As a consequence of defective hemoglobin processing, *Pf*ATG18-deficient parasites produced only 62% of physiological hemozoin levels by 36 hpi, as determined by quantitative polarization microscopy (Figure 3K). Together, these findings suggest that while *Pf*ATG18-deficient parasites remain capable of endocytosing host cell cytosol, they ultimately fail to deliver the cargo to the DV.

### DV fragmentation precedes HTC accumulation in *Pf*ATG18-deficient parasites

We next aimed to dissect the sequence of events underlying the dual phenotype of *Pf*ATG18-deficient parasites. To determine whether early DV fragments remain capable of receiving cargo, we treated 28-hour old *atg18:loxP + crt-mNG* parasites with E64. This broad-spectrum protease inhibitor blocks hemoglobin degradation and induces DV swelling due to accumulation of undigested hemoglobin, unless endocytic trafficking is impaired (*51, 52*). Live microscopy of CRT-mNG revealed that 8 hours of E64 treatment caused significant vacuolar bloating in DMSO control parasites (Figure 4A). In RAP-treated parasites, we observed an E64-dependent increase in the size of individual DV fragments. Because the DV of these parasites was already fragmented at the time of E64 addition, these results indicate that early DV fragments had received endocytic cargo during the 8-hour treatment window (Figure 4A). This was further supported by microscopic analysis of Giemsa-stained blood smears, which revealed host cell cytosol accumulated in bloated DV fragments of E64-treated *Pf*ATG18-null parasites (Figure 4B).

**Figure 4.**
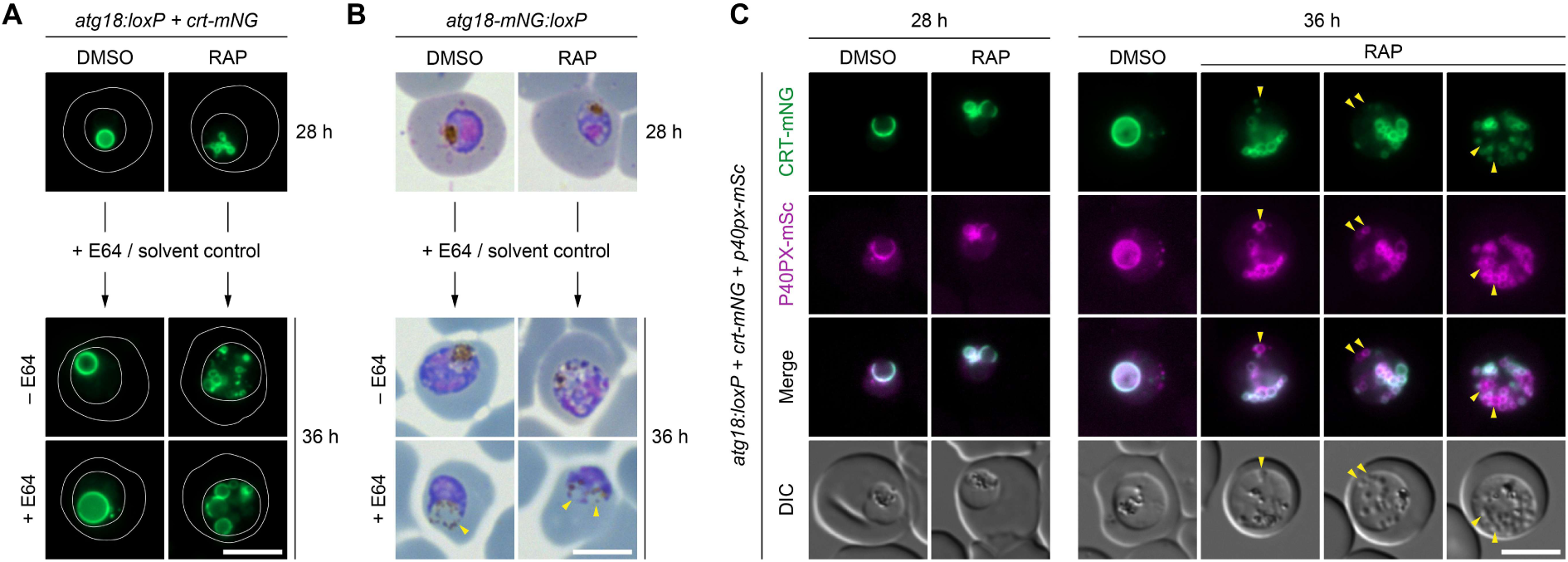
Vacuolar fission precedes HTC accumulation in *Pf*ATG18-deficient parasites. Synchronized parasites were treated with DMSO or RAP from the ring stage onward and imaged at 28 and 36 hpi. **(A,B)** Early DV fragments in *Pf*ATG18-deficient parasites still receive cargo. E64-mediated inhibition of hemoglobin degradation causes bloating in both intact and fragmented DVs. Representative images of CRT-mNG (A) and Giemsa-stained blood smears (B). White outlines: parasite and host cell; yellow arrowheads: bloated DV (fragments) containing host cell cytosol and hemozoin. **(C)** In *Pf*ATG18-deficient parasites, the PI3P-marker P40PX-mSc remains restricted to the DVM at the onset of fragmentation and only later appears on accumulating HTCs. Yellow arrowheads: P40PX-mSc-positive HTCs lacking CRT-mNG labelling. Vesicular compartments are visible in DIC at 36 hpi, but not at 28 hpi. Scale bars: 5 µm.

To allow for the simultaneous monitoring of DV morphology and endocytic hemoglobin delivery, we introduced into the *atg18:loxP + crt-mNG* parasites an expression cassette for P40PX-mSc, a PI3P-binding fluorescent reporter that marks both the DVM and HTCs (*24, 53*). We imaged these transgenic parasites by live fluorescence microscopy at 28 and 36 hpi. At both timepoints, DMSO-treated control parasites exhibited an intact DV, with P40PX-mSc localizing almost entirely to the vacuolar membrane, indicative of efficient PI3P transfer from HTCs to the DV (Figure 4C). Although RAP-treated parasites exhibited a fragmented DV already at 28 hpi, P40PX-mSc was found associated exclusively with CRT at that time. This changed dramatically 8 hours later, when a substantial fraction of P40PX-mSc localized to vesicular compartments devoid of CRT, identifying them as HTCs (Figure 4C). Together, these findings suggest that in *Pf*ATG18-deficient parasites, vacuolar fragmentation precedes HTC accumulation.

### Loss of *Pf*ATG18 hypersensitizes ring-stage parasites to dihydroartemisinin

Perturbations in hemoglobin handling and DV physiology can both confer resistance to drugs whose activity depends on hemoglobin consumption, such as chloroquine and artemisinin (*15–18*). Reports linking a T38I mutation in *Pf*ATG18 to prolonged parasite clearance half-life in artemisinin-treated patients (*45, 46*) therefore motivated us to examine whether *Pf*ATG18-deficient parasites might similarly exhibit reduced drug sensitivity.

To test this, we first established conditions that allowed us to quantify parasite replication as a readout in drug sensitivity assays. We delayed RAP induction so parasites could produce minimal amounts of *Pf*ATG18 and complete one replication cycle. When RAP was applied at 16 hpi, parasites progressed into the second intraerythrocytic cycle at ∼37% efficiency, compared to DMSO controls (Figure 5A). At this induction time, *Pf*ATG18 protein levels were reduced by ∼90% at the end of the first cycle (Figure 5B). Live-cell imaging revealed a severely fragmented DV without significant accumulation of HTCs, indicating that *Pf*ATG18-dependent functions were disrupted in schizonts, although the delayed induction bypassed the fusion defect during the first cycle (Figure 5C).

**Figure 5.**
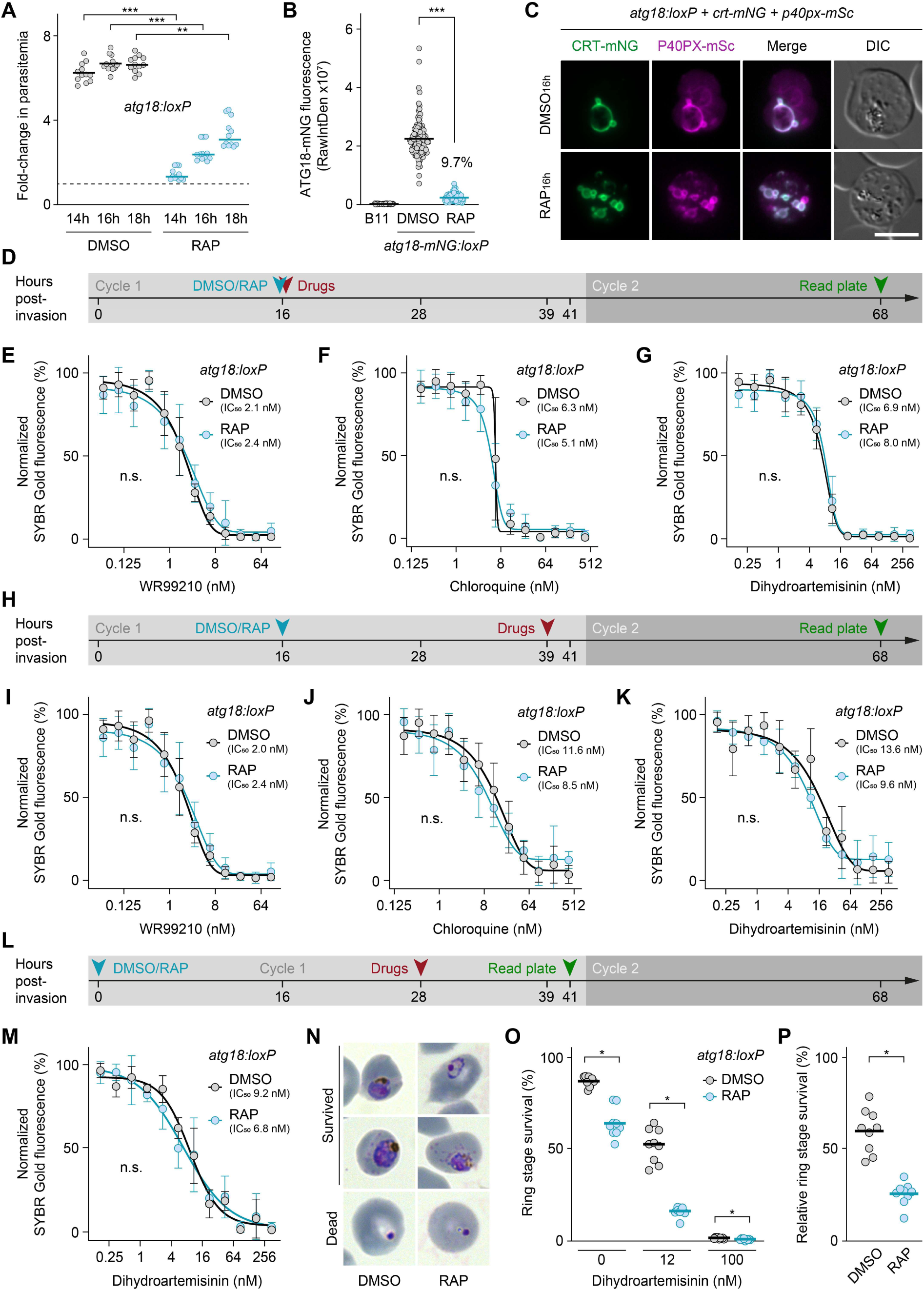
Drug sensitivity profiling of *Pf*ATG18-deficient parasites. **(A)** Delayed inactivation of *Pf*ATG18 allows completion of one intraerythrocytic cycle. Parasitemia was measured after invasion and again at 68 hpi, and fold-change was calculated. DMSO or RAP was added at the indicated time points. Dashed line: no change. Individual and mean values; paired *t*-test; n = 4 experiments with 3 technical replicates each. **, P < 0.01; ***, P < 0.001. **(B,C)** Live fluorescence microscopy shows that RAP induction at 16 hpi strongly depletes *Pf*ATG18 (B) and causes DV fragmentation in schizonts (C). Individual and mean values; paired *t*-test; n = 132 parasites, 4 experiments. **(D-G)** Drug sensitivity assays with induction and drug addition at 16 hpi and readout in the second cycle (D) show that *Pf*ATG18-deficient parasites retain normal sensitivity to WR99210 (E), chloroquine (F), and DHA (G). **(H-K)** Delaying drug addition to schizont stage (H) similarly shows no change in drug sensitivity (I-K). **(L,M)** Drug sensitivity assays with ring-stage induction, drug addition at DV fragmentation onset, and readout at the end of the first cycle (L) reveal no change in DHA sensitivity upon *Pf*ATG18 loss (M). IC_50_ values were determined by non-linear regression. Mean +/− SEM; n = 3 experiments with 3 technical replicates each. n.s., P > 0.05. **(N-P)** *Pf*ATG18 loss induced at 16 hpi increases DHA sensitivity in second-cycle ring stages. (N) Representative parasite morphologies, showing surviving and dead forms. (O) Ring-stage survival in response to a 6-hour DHA pulse at indicated concentrations. (P) Relative ring-stage survival after a 12 nM DHA pulse, normalized to baseline survival in untreated controls. Individual and mean values; paired *t*-test; n = 3 experiments with 3 technical replicates each. *, P < 0.05. Scale bars: 5 µm.

Using this setup, we assessed drug sensitivity by exposing parasites at the time of induction to increasing concentrations of dihydroartemisinin (DHA), chloroquine, or the negative control compound WR99210, which inhibits folate biosynthesis (Figure 5D). Parasite DNA was quantified 68 hpi using SYBR Gold (*54*). The resulting growth-response curves showed no RAP-dependent shifts in IC_50_ values for any of the drugs (Figure 5E-G). To address the possibility that drug action might have occurred before loss of *Pf*ATG18 function, we repeated the assays with drug addition late in schizogony (Figure 5H). Again, no differences in drug-sensitivity were observed (Figure 5I-K).

To examine whether the loss of HTC-DV fusion (bypassed under the previous induction schedule) could confer drug resistance, we treated cultures with DMSO or RAP immediately after invasion and added DHA 28 hours later (Figure 5L). At that time, the DV was already fragmented and HTC accumulation was imminent (Figure 4). Quantification of replicated DNA at the end of the first cycle yielded robust drug-response curves, yet again showing no differences in DHA sensitivity (Figure 5M).

Because artemisinin resistance is associated with increased ring-stage survival (*55, 56*), we next induced *Pf*ATG18 loss at 16 hpi and treated second-cycle rings with DHA for 6 hours, then assessed their survival. In DHA-free controls, DMSO-treated parasites matured normally into healthy second-cycle trophozoites, whereas RAP-treated parasites arrested as late rings or dysmorphic young trophozoites, with some parasites appearing dead, indicating that delayed *Pf*ATG18 loss already impairs second-cycle development (Figure 5N). A 100 nM DHA pulse caused near-complete death in both conditions, producing uniformly pyknotic forms that were readily distinguishable from surviving parasites (Figure 5N,O). A 12 nM DHA pulse reduced survival from 87% to 51% in DMSO-treated parasites and from 64% to 16% in RAP-treated parasites (Figure 5O). After accounting for baseline survival differences, *Pf*ATG18-deficient parasites were approximately twice as sensitive to DHA at this stage (Figure 5P).

Together, these data demonstrate that loss of *Pf*ATG18 alone does not confer resistance to chloroquine or DHA. Rather, it renders young parasite stages more vulnerable to chemotherapeutic insult.

### Loss of *Pf*ATG18 results in V-ATPase disassembly

We recently reported that DVM integrity depends on the parasite’s V-ATPase complex (*20*) (Figure 6A). Genetic inactivation of individual V_0_ sector subunits caused disassembly of the proton pump and triggered vacuolar fission – an effect similar to that observed in *Pf*ATG18-deficient parasites (*20*). This raises the possibility that *Pf*ATG18 may play a role in regulating V-ATPase stability and function.

**Figure 6.**
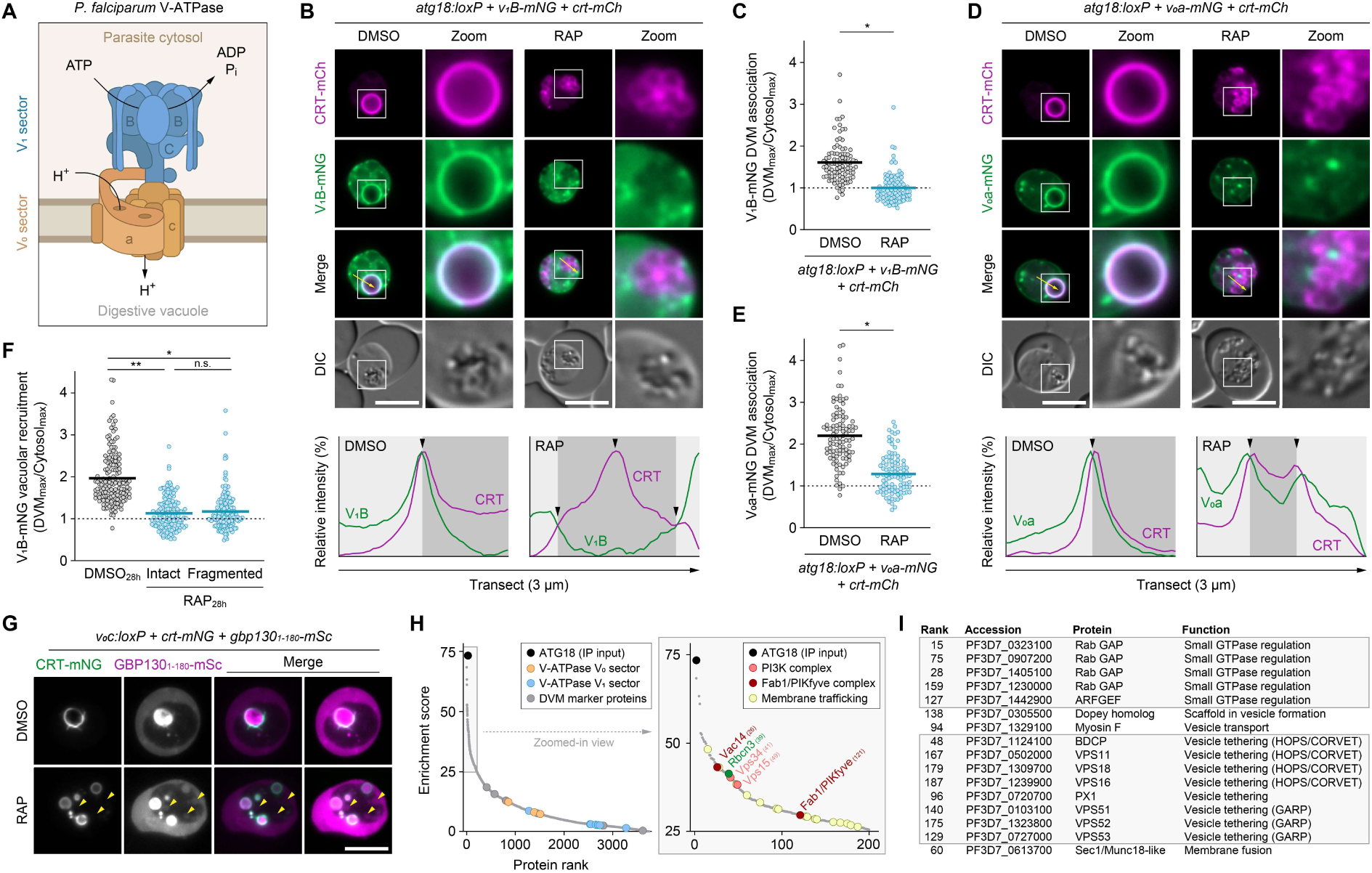
*Pf*ATG18 is critical for V-ATPase integrity. **(A)** Schematic of the V-ATPase complex, with sectors and relevant subunits indicated. **(B,C)** Subunit V_1_B dissociates from the DVM upon *Pf*ATG18 loss. (B) Live colocalization of CRT-mCh and V_1_B-mNG. Yellow arrows: transects for intensity plots; light gray: parasite cytoplasm; dark gray: DV; black arrowheads: DVM. (C) Ratio of DVM– to cytosol-localized V_1_B-mNG intensity. Individual and mean values; paired *t*-test; n = 99 parasites, 3 experiments. *, P < 0.05. Parasites were induced at ring stage and imaged at 36 hpi. The dashed line denotes full equilibration of both V_1_B pools. **(D,E)** Subunit V_0_a is strongly depleted from the DVM in *Pf*ATG18-deficient parasites. Analysis as in (B,C). **(F)** V_1_B dissociation precedes DV fragmentation. Loss of *Pf*ATG18 was induced at 28 hpi, shifting the onset of DV fragmentation to 36 hpi. Analysis as in (C). Individual and mean values; one-way ANOVA with Tukey’s test; n = 165 parasites, 5 experiments. n.s., P > 0.05; **, P < 0.01. **(G)** Conditional inactivation of subunit V_0_c recapitulates the *Pf*ATG18 knockout phenotype, with DV fission and HTC accumulation (arrowheads). Shown are CRT-mNG and GBP130_1-180_-mSc, an exported and reinternalized marker. **(H,I)** Co-immunoprecipitation combined with mass spectrometry identifies candidate *Pf*ATG18 interaction partners involved in phosphoinositide balance and membrane trafficking. (H) Protein hits ranked by enrichment score, defined as the log_2_ fold-change (*atg18-mNG:loxP* / B11) multiplied by the negative log_10_ of the p-value. Benjamini-Hochberg corrected *t*-test; n = 3 experiments with 3 technical replicates each. Relevant proteins as well as unrelated DVM markers are indicated. (I) Membrane trafficking-related proteins among the top 200 hits with a confirmed or potential role in the endolysosomal pathway. Scale bars: 5 µm.

To investigate this, we first examined the localization of mNG-tagged subunit V_1_B in *atg18:loxP* parasites. In DMSO-treated controls, V_1_B localized predominantly to the DVM surrounding the hemozoin crystals. Upon RAP treatment, however, V_1_B no longer associated with the hemozoin clusters and instead localized diffusely throughout the cell, indicating dissociation from the DVM (Supplemental Figure S7A). A similar pattern was observed for subunit V_1_C, suggesting that *Pf*ATG18 loss broadly disrupts membrane association across the entire V_1_ sector (Supplemental Figure S7B). Because fragmentation hindered reliable identification of the DVM, we further modified the *atg18:loxP + v_1_B-mNG* line to express mCh-tagged CRT. Following *Pf*ATG18 inactivation, V_1_B no longer colocalized with CRT at the fragmented DVM (Figure 6B), and quantification of membrane-associated versus cytosolic signals revealed that these V_1_B pools had fully equilibrated in *Pf*ATG18-deficient parasites (Figure 6C).

The dissociation of V_1_ sector subunits indicates defective V-ATPase assembly and has been shown to occur in response to perturbations of either the V_0_ or the V_1_ sector (*20*). Importantly, we have shown that DV fragmentation is exclusively associated with V_0_ sector dysfunction, whereas perturbations of the V_1_ sector alone do not produce this effect (*20*). We therefore investigated the localization of subunit V_0_a in *Pf*ATG18-deficient parasites and found it to be severely depleted from the vacuolar membrane (Supplemental Figure S7C). As with V_1_B, we introduced CRT-mCh as a DVM marker and confirmed that V_0_a association with CRT-defined DV fragments was markedly reduced, although dissociation was not complete (Figure 6D,E).

Our findings suggest that V-ATPase disassembly may be the proximal cause of DV fragmentation in *Pf*ATG18-deficient parasites. To determine whether disassembly precedes fragmentation, we treated the *atg18:loxP + v_1_B-mNG + crt-mCh* line with RAP at 28 hpi and analyzed V_1_B localization eight hours later, at the onset of DVM fission. At this stage, approximately one half of the parasite population retained an intact DV, while the other half had initiated vacuolar fragmentation. In marked contrast to DMSO controls, V_1_B was almost completely dissociated from the DVM in RAP-treated parasites, regardless of whether the DV was already fragmented or not, indicating that *Pf*ATG18 inactivation disrupts V-ATPase assembly before vacuolar integrity is compromised (Figure 6F, Supplemental Figure S7D).

To determine whether V-ATPase dysfunction could explain both vacuolar fragmentation and the fusion defect in *Pf*ATG18-deficient parasites, we tested whether directly compromising V_0_ sector integrity recapitulates this dual phenotype. We modified a previously generated conditional knockout line for subunit V_0_c expressing CRT-mNG (*20*) to additionally express the exported host cell cytosol reporter GBP130_1-180_-mSc. As previously shown, V_0_c inactivation caused fission of the DV into numerous CRT-positive compartments (*20*), which contained high levels of GBP130_1-180_-mSc (Figure 6G). Importantly, the host cytosolic marker was also detected in accumulating spherical compartments lacking CRT, at concentrations comparable to the erythrocyte cytosol, suggesting that they represent hemoglobin trafficking intermediates upstream of the DV (Figure 6G). These findings indicate that inactivation of V_0_ sector subunits affects both vacuolar integrity and fusion. Lower abundance of fully assembled V_0_ sectors at the DVM could thus sufficiently explain the dual defects observed in *Pf*ATG18-deficient parasites.

### *Pf*ATG18 associates with a putative V-ATPase assembly factor, phosphoinositide-converting enzymes and vesicle trafficking machinery

We next sought to determine whether *Pf*ATG18 influences V-ATPase function through direct physical interactions or indirectly. To identify potential *Pf*ATG18 interaction partners, we immunoprecipitated the protein from *atg18-mNG:loxP* parasites and analyzed co-purifying proteins by mass spectrometry (Supplemental Dataset 1). After ranking hits by enrichment and statistical significance relative to parental controls, we found that none of the V-ATPase subunits ranked highly (Figure 6H), suggesting that the contribution of *Pf*ATG18 to V-ATPase assembly is unlikely to involve direct contact with the proton pump.

Instead, we detected among the highest ranking hits a conserved parasite protein of 721 kDa annotated as a Rbcn3 homolog, which in other eukaryotes acts as a regulatory scaffold mediating V_0_-V_1_ coupling (*36*). Other top-ranking proteins included two components each of the PI3K and Fab1/PIKfyve complexes, which synthesize PI3P and PI(3,5)P_2_, respectively (Figure 6H), as well as multiple membrane trafficking factors, including a Sec1/Munc18-like protein and several small GTPase regulators (Figure 6H,I). Four components of the HOPS/CORVET vesicle-tethering machinery, the phosphoinositide-binding protein PX1, and Myosin F were also enriched – all previously implicated in hemoglobin trafficking (*23, 26, 57*). By contrast, components of the retromer complex, which have been reported to interact with ATG18 in yeast and mammalian cells, ranked low in our analysis, consistent with the observed lack of colocalization (Figure 1I,J).

Although not all enriched proteins may represent direct *Pf*ATG18 interactors, our findings suggest that *Pf*ATG18 might function as a versatile interaction hub at the DVM. It may recruit, stabilize, or regulate diverse proteins and assemblies that support core DV functions, including membrane fusion, lipid conversion, and V-ATPase assembly, thereby exerting broad effects on organelle physiology.

## DISCUSSION

Malaria parasites possess only a limited subset of autophagy-related genes and lack key components required for autophagy initiation and autophagosome membrane expansion (*43, 58*). Beyond *Pf*ATG18 and PI3K, they retain an ortholog of ATG8, along with the machinery necessary for its processing and membrane conjugation. In classical autophagy, ATG8 functions as a central hub, orchestrating autophagosome formation, cargo recognition, and membrane dynamics (*59*). In *P. falciparum*, however, *Pf*ATG8 localizes specifically to the apicoplast, and its absence results in lethal defects in organelle inheritance (*60, 61*). Remarkably, survival of *Pf*ATG8-deficient parasites can be fully restored by supplying the apicoplast-derived metabolite isopentenyl pyrophosphate, thus bypassing the organelle’s essential metabolic functions (*61*). These findings, alongside a growing body of evidence, indicate that malaria parasites have lost many components of the autophagic machinery during evolution, while others have been repurposed for roles beyond classical autophagy. Similarly, non-canonical functions have been proposed for ATG18 in apicomplexan parasites, but also in other model organisms, including roles in apicoplast inheritance (*44*), retromer engagement (*40, 62*), and endocytic membrane fission (*39, 63*). In our study, however, neither localization, phenotype, nor protein interaction data provided evidence for an involvement of *Pf*ATG18 in apicoplast– or retromer-related processes.

Instead, we found that loss of *Pf*ATG18 leads to multiple cellular abnormalities, all linked to the parasite’s endocytic pathway. The most striking of these were fragmentation of the DV and accumulation of HTCs, together indicating that *Pf*ATG18 directs endocytic membranes toward fusion rather than fission. Accordingly, *Pf*ATG18 may facilitate both heterotypic fusion between the DV and HTCs and homotypic fusion between the DV and its lobes. Our findings contrast with reports from yeast and mammalian cells, where ATG18 homologs function in membrane scission: disruption of these proteins prevents vacuoles or endosomes from undergoing fission, causing organelle enlargement (*40, 63*). Although DV fragmentation is the exact opposite of these phenotypes, *Pf*ATG18 expression was shown to rescue the vacuolar fission defect in ATG18-deficient yeast, highlighting how conserved molecular functions can produce different outcomes depending on cellular context (*37*).

In *Pf*ATG18-null parasites, HTCs accumulate only after DV fragmentation has already commenced, suggesting that their failure to fuse could be a secondary consequence of a disrupted DV. However, this interpretation is challenged by two observations: (1) early DV fragments can still receive cargo, and (2) other *Plasmodium* species naturally maintain multiple small DVs yet process hemoglobin just as efficiently (*64, 65*). An alternative possibility is that *Pf*ATG18 regulates DV maintenance and fusion through distinct pathways by serving as a scaffold for multiple molecular assemblies at the DVM, a functional versatility commonly observed in WD40 repeat proteins (*66*). This notion is consistent with the identification of functionally distinct protein assemblies as potential *Pf*ATG18 interactors, including vesicle tethering complexes, phosphoinositide-converting assemblies, and other protein machinery involved in endocytic trafficking. While our Co-IP data identify strong candidates for *Pf*ATG18 binding, complementary proximity labeling studies may help determine whether these interactions also occur in the context of the intact parasite cell.

Our observation that the V-ATPase disassembles in response to *Pf*ATG18 inactivation raises the possibility that the defects in endocytic membrane dynamics may arise directly from proton pump dysfunction. The V-ATPase has been implicated in membrane fusion and fission across different organisms, although it often remains unclear whether these effects are structural or pH-dependent (*67, 68*). In yeast, vacuole fusion is facilitated by interactions between opposing V_0_ sectors in *trans*, promoting lipid mixing and hemifusion, which likely involves V_0_-SNARE interactions (*69, 70*). Salt-induced vacuole fission has also been reported to depend on V-ATPase activity (*71*). We show that in *P. falciparum*, targeted interference with V_0_ sector integrity phenocopies the dual defects observed in *Pf*ATG18 knockout parasites and that disruption of *Pf*ATG18 is accompanied by V_0_ sector destabilization, leading to complete loss of V_0_-V_1_ coupling. Our findings may also provide a mechanistic explanation for previous observations in yeast, where ATG18 was reported to associate cyclically with the vacuole in strict synchrony with organellar pH oscillations (*72*), and to ensure vacuolar pH asymmetry during budding (*73*).

We found no evidence for direct interactions between *Pf*ATG18 and the V-ATPase. Instead, among the highest-ranking hits in *Pf*ATG18 pull-downs was a putative Rbcn3 homolog. In other eukaryotes, Rbcn3 – or its yeast analog, the RAVE complex – facilitates V_0_-V_1_ association, particularly when the two sectors have dissociated in response to cellular stress signals (*36*). While V_1_ dissociation in *Pf*ATG18-null parasites may result from loss of Rbcn3 function, the destabilization of the V_0_ sector is more difficult to reconcile with this mechanism, unless *Plasmodium* Rbcn3 has evolved additional roles in V-ATPase assembly beyond sector coupling. It thus appears likely that *Pf*ATG18 may influence the proton pump’s membrane sector indirectly by modulating conditions at the DVM that support its stability and function.

One such condition could be the homeostasis of phosphoinositide lipids. Indeed, multiple components of the PI3K and Fab1/PIKfyve complexes ranked among the top hits in *Pf*ATG18 pull-downs. In yeast, PI(3,5)P_2_ produced by Fab1 directly binds to the N-terminal domain of V_0_a which helps anchor the subunit in the membrane and stabilizes V_0_-V_1_ interactions (*74, 75*). Similarly, disruption of PI3K-mediated PI3P synthesis, resulted in significantly reduced levels of both V_0_ and V_1_ sector subunits in yeast vacuolar vesicle preparations (*76*). It is therefore tempting to speculate that the phenotypes observed in our mutant may largely result from disturbed phosphoinositide homeostasis, with *Pf*ATG18 regulating their balance at the DVM by modulating PI3K and Fab1/PIKfyve activity. In yeast, ATG18 has been proposed to act as a negative regulator of PI(3,5)P_2_ levels. In this model, ATG18 is recruited from the cytosol to the vacuole membrane when PI(3,5)P_2_ levels spike, often in response to osmotic stress. Elevated PI(3,5)P_2_ would then promote ATG18 binding to the membrane, where it could downregulate Fab1/PIKfyve activity and limit further PI(3,5)P_2_ production (*77*). In contrast to yeast, *Pf*ATG18 does not appear to exist in a cytosolic pool, arguing against a recruitment-based regulation mechanism and rather supporting a constitutive role at the DVM.

While PI3P is highly enriched at the DVM, PI(3,5)P_2_ has not yet been detected in the parasite, either due to technical limitations arising from its low abundance, or because it may be absent entirely (*34*). In support of the latter scenario, *Pf*ATG18 interacted only with PI3P in liposome binding assays, whereas its *T. gondii* ortholog bound both PI3P and PI(3,5)P_2_ (*44*), although broader phosphoinositide interactions were observed in protein-lipid overlay assays (*45*). Remarkably, when expressed in ATG18-deficient yeast, *Pf*ATG18 reestablishes vacuolar fission, a process previously shown to depend on PI(3,5)P_2_ (*32, 37*). Thus, even if the *Plasmodium* Fab1/PIKfyve complex has lost its catalytic activity during evolution, its interaction with ATG18 may still be conserved and mediate functions beyond phosphoinositide balance. Mechanistic studies on the parasite’s Fab1/PIKfyve complex, combined with sensitive phosphoinositide biosensors, are thus needed to clarify the role of PI(3,5)P_2_ in parasite development and its relationship with *Pf*ATG18.

Our SBF-SEM analysis allowed three-dimensional reconstruction of accumulating HTCs, revealing distinct membrane profiles closely associated with the HTC envelope. These slit-like invaginations frequently terminated in a small intralumenal vesicles whose electron density suggests they contain host cell cytosol. It remains unclear whether these features represent fission scars from HTC budding at the cytostome, or form through membrane inward bending afterwards. The latter scenario would be consistent with our observation that the slits often originate from funnel-like pits, reminiscent of ESCRT-mediated intraluminal vesicle formation (*78*). Vesicular HTC subcompartments have been observed previously by conventional transmission electron microscopy independently of *Pf*ATG18 and therefore appear to represent a general feature of HTC physiology (*29, 51, 79, 80*). Our SBF-SEM dataset provides the first three-dimensional characterization of their ultrastructure; however, higher-resolution approaches such as cryogenic electron tomography will be required to further resolve their organization and membrane architecture.

The role of *Pf*ATG18 in hemoglobin trafficking offered a plausible mechanistic framework for the association between the T38I point mutation and prolonged parasite clearance in artemisinin-treated patients (*45, 46*). We hypothesized that reduced hemoglobin delivery, and consequently diminished heme release in the DV, would decrease artemisinin sensitivity in *Pf*ATG18-deficient parasites due to reduced drug activation (*55, 81*). However, despite multiple assay variations, including optimized induction and drug incubation schedules, we were unable to support this hypothesis. This is consistent with a previous study examining the isolated effect of the T38I mutation, which reported no changes in sensitivity to artemisinin-based antimalarials, using both standard growth-inhibition and ring-stage survival assays (*45*). It therefore remains possible that resistance effects conferred by the T38I mutation are too subtle to resolve under laboratory conditions but may nonetheless provide a selective advantage in the field. Alternatively, the T38I mutation may confer a selective advantage only in specific genetic backgrounds that harbor additional mutations linked to artemisinin resistance.

To our surprise, loss of *Pf*ATG18 rendered ring-stage parasites hypersensitive to DHA. This hypersensitivity could arise for different reasons, the simplest being that parasites already compromised in viability are generally more susceptible to chemotherapeutic insult. Alternatively, the phenotype could reflect disturbed phosphoinositide homeostasis, as elevated levels of PI3P have previously been linked to artemisinin resistance (*28, 82*). If *Pf*ATG18-deficient parasites exhibit a phosphoinositide balance shifted toward lower PI3P and higher PI(3,5)P_2_, like their yeast counterparts (*77*), this could explain the observed sensitivity profile. However, a mechanistic link between *Pf*ATG18, phosphoinositide homeostasis, and artemisinin resistance remains to be established in future studies.

## METHODS

### Structural homology modelling

An AlphaFold3 model of *Pf*ATG18 was aligned to the crystal structure of the *Sc*ATG18 β-propeller core (*38*). Structural alignment was performed in PyMol (version 2.5.7, Schrödinger, LLC) using the “cealign” command, and the RMSD value was calculated for the aligned residues. Due to low-confidence predictions, the 4CD and 6CD loops extending from the *Pf*ATG18 β-propeller core were excluded from the alignment.

### Parasite cultivation

*Plasmodium falciparum* parasites were maintained at 37°C in human erythrocytes (B+) suspended in Roswell Park Memorial Institute medium 1640 (RPMI 1640) supplemented with 0.5% AlbuMax II. Cultures were kept in an atmosphere of 1% O_2_, 5% CO_2_, and 94% N_2_, and synchronized as previously described (*20*). Briefly, mature schizonts were isolated by centrifugation over a 60% Percoll cushion and allowed to rupture and invade fresh erythrocytes for 2 hours under constant agitation. Residual schizonts were subsequently removed using a second Percoll gradient, followed by 5% sorbitol treatment of the infected cells, yielding highly synchronous ring-stage cultures. Parasitemia was determined either by microscopy of Giemsa-stained blood smears or by flow cytometry of SBYR Green-stained cultures and adjusted by dilution with uninfected erythrocytes. Conditional *Pf*ATG18 knockout lines were treated with DMSO or RAP from the ring stage onward, followed by phenotypic analysis at 36 hpi, unless otherwise specified. The *v_0_c:loxP + crt-mNG + gbp130_1-180_-mSc* parasites were induced and analyzed as described previously (*20*). E64 and 5-aminolevulinic acid were added to the culture medium at 28 hpi at final concentrations of 21.7 µM or 200 mM, respectively.

### Generation and validation of transgenic parasites

The *atg18:loxP* parasite line was generated via Cas9-guided homologous integration of a synthetic repair template containing recodonized *Pf*ATG18, flanked by two *loxP* sites – one within an artificial intron and the other immediately downstream of the stop codon (*48*) (Figure 1B). In the *atg18-mNG:loxP* line, the C-terminus was additionally tagged with mNG. These genetic modifications were introduced into the 3D7-based B11 line, which expresses dimerizable Cre recombinase (DiCre) and permits RAP-induced excision of *loxP*-flanked genomic sequences (*47*).

For Cas9-mediated genomic integration, ∼10^8^ Percoll-enriched schizonts were electroporated using a Lonza Amaxa 4D Nucleofector device, introducing 20 µg of guide plasmid and 60 µg of linearized repair template. Following transfection, parasites were allowed to invade fresh erythrocytes, and drug selection with 2.5 nM WR99210 was applied 24 hours later for a total duration of 4 days. Transgenic parasite clones were isolated 1-2 weeks later using a previously described plaque assay (*83*). Correct genomic integration of the repair templates and RAP-induced excision of *loxP*-flanked sequences were verified by diagnostic PCR using primer combinations shown in Figure 1B and Supplemental Table S1.

Endogenous tagging of CRT, PM2, HRP2, FT2, Vps26, Vps29, V_1_B, V_1_C, and V_0_a was also achieved by Cas9-mediated genome editing. Clonality of the resulting mutants was confirmed by live fluorescence microscopy. mSc-Rab5a (*26*), mSc-Rab7 (*26*), P40PX-mSc (*51*), and GBP130_1-180_-mSc (*84*) fusion proteins were expressed episomally under constant drug pressure with 0.9 µM DSM1. Repair templates and Cas9 guide sequences are provided in Supplemental Tables S2 and S3, respectively. All transgenic parasite lines used in this study are listed in Supplemental Table S4.

### Analysis of parasite replication and drug sensitivity

To analyze parasite replication, ring-stage cultures were adjusted to an initial parasitemia of 0.1%, and parasitemia was monitored over four intraerythrocytic cycles at two-day intervals. The percentage of infected erythrocytes was determined by flow cytometry of cultures stained for 40 minutes at 37°C in medium containing 1:10,000 SYBR Green, followed by 40 minutes of de-staining in plain medium. Samples were analyzed on an automated Agilent NovoCyte 3000VYB flow cytometer equipped with a NovoSampler Pro.

To assess parasite drug sensitivity, synchronized *atg18:loxP* cultures were adjusted to 2% parasitemia at 2% hematocrit and incubated in 96-well black-bottom plates with DMSO/RAP and varying drug concentrations in a final volume of 200 µL. Cells were lysed at 68 hpi, and DNA was stained with SYBR Gold, as described previously (*54*). Fluorescence was quantified using a Tecan GENios plate reader, with excitation and emission wavelengths set to 485 nm and 535 nm, respectively. Raw fluorescence values were corrected for non-infected erythrocyte controls and normalized to the highest and lowest readings. IC_50_ values were calculated using non-linear regression in GraphPad Prism (version 10.4.1), applying the “log(inhibitor) vs normalized response – Variable slope” equation.

To assess ring-stage survival following DHA treatment, *atg18:loxP* parasites were induced at 16 hpi. First-cycle schizonts were isolated and added to fresh erythrocytes to generate tightly synchronized second-cycle ring stages at ∼5% parasitemia and 2% hematocrit, seeded into 24-well plates at a total volume of 600 µL per well. Ring-stage parasites were then exposed to 0, 12, or 100 nM DHA for 6 hours at 37°C. Following the drug pulse, cultures were washed three times with medium, resuspended in 600 µL per well, and incubated at 37°C. Parasite survival was assessed 18 h later by microscopic examination of Giemsa-stained blood smears, with viable and pyknotic parasites quantified by manual counting.

### Light and fluorescence microscopy

Light and fluorescence microscopy were performed using a DM6 B microscope equipped with a K8 B/W camera and a K3C color camera, all from Leica. The microscope was fitted with two crossed polarizers, enabling *in situ* visualization and quantification of birefringent hemozoin crystals. All images were analyzed and processed using FIJI (version 2.16.0/1.54p). Transmitted polarized light and fluorescence signals were measured as mean intensity, maximum intensity or raw integrated density, with appropriate background corrections applied in all cases.

All fluorescence imaging was performed with live parasites. To visualize dimmer pools of PPIX, HRP2-mCh, or GBP130_1-180_-mSc, selected micrographs were intentionally oversaturated. Signals specific to CRT-mCh or ATG18-mNG in the respective double mutant were identified by first normalizing the input signals to their respective minimum and maximum pixel values. The normalized intensities were then subtracted on a pixel-by-pixel basis, and the resulting differences were visualized as pseudo-images. To quantify the membrane association of V_1_B-mNG and V_0_a-mNG, fluorescence intensity was measured in randomly selected regions of the DVM and parasite cytoplasm, identified by the presence or absence of CRT-mCh labelling, respectively. Ratios of the maximum intensity values were calculated.

Parasite size was determined microscopically in Giemsa-stained blood smear as the percentage of the erythrocyte area occupied. To quantify the hemozoin dispersal area, the polygon selection tool in FIJI was used to manually outline a single region per parasite that enclosed all hemozoin particles, minimizing the number of nodes and avoiding acute polygon angles. The resulting dispersal area was then normalized to parasite size.

### Serial block-face scanning electron microscopy

For SBF-SEM analysis, infected erythrocytes were enriched 36 hpi via Percoll gradient centrifugation. Cells were fixed in 2% paraformaldehyde and 2.5% glutaraldehyde in PBS, then embedded in 6% gelatin. Post-fixation was performed with 2.5% glutaraldehyde in PBS, followed by staining with 2% OsO_4_, 1.5% potassium ferrocyanide and 2 mM CaCl_2_ in PBS on ice. Samples were then sequentially incubated with 0.5% thiocarbohydrazide, 2% OsO_4_ and 0.1% gallic acid. Subsequent overnight staining was performed at 4°C with 1% uranyl acetate. The following day, samples were stained with Walton’s lead aspartate at 60°C and dehydrated through a graded ethanol series on ice.

Samples were infiltrated with 50% and 70% Epon in ethanol, followed by two incubations in 100% Epon and polymerization with 3% silver flakes at 60°C. Sample blocks measuring 0.5 x 0.5 mm were cut, mounted, and placed into a Gatan 3View stage within a Jeol JSM-7100F scanning electron microscope. Imaging was performed at 3 kV acceleration voltage with a stage bias of +500 V, using a probe current setting of 1, 3 nm pixel size, and 50 nm section thickness. Acquisition and stack alignment were carried out with Gatan Digital Micrograph (version 3.32). Images were processed in FIJI using the StackReg plugin for alignment and the DenoisEM plugin (*85*) with Tikhonov algorithm for denoising. Segmentation and 3D reconstruction were performed using Microscopy Image Browser (*86*) (version 2.9.1) and Imaris (version 9.8.2, Oxford Instruments).

### Western blotting

For Western blot analysis, parasites were released from host erythrocytes by treatment with 0.03% saponin in PBS for 10 minutes on ice. The parasites were washed several times with PBS until the supernatant was colorless, and then lysed in PBS containing 4% SDS, 0.5% Triton X-100, and a protease inhibitor cocktail. Lysates were mixed with Laemmli buffer and incubated at 95°C for 5 minutes prior to SDS-PAGE.

Separated proteins were either stained in-gel using Coomassie Brilliant Blue R250 or transferred onto nitrocellulose membranes. Primary antibodies were used at the following concentrations: α-BiP ((*87*); rabbit; 1:2,000), α-mNG (ChromoTek; mouse; 1:1,000), α-Hb (Abcam; rabbit; 1:2,000). Bound primary antibodies were detected by chemiluminescence using horseradish-peroxidase-conjugated secondary antibodies (Agilent; 1:5,000 – 1:3,000) and ECL reagent.

### Co-immunoprecipitation

To identify potential *Pf*ATG18-interacting proteins, *Pf*ATG18 was immunoprecipitated from *atg18-mNG:loxP* parasites using the mNeonGreen-Trap Magnetic Agarose kit (ChromoTek). Briefly, 200-mL cultures at 7% parasitemia and 1% hematocrit were harvested, and parasites were released by treatment with 0.03% saponin. Following centrifugation, parasite pellets were washed with PBS and used as input material for Co-IP. The protocol was performed according to the manufacturer’s instructions with minor modifications. For each Co-IP experiment, pellets were lysed in 800 µL of NP40-based lysis buffer, after which the lysate was diluted 1:2 with dilution buffer and incubated overnight at 4°C with anti-mNG nanobody-coupled magnetic beads under constant agitation. The beads were then washed four times and heated at 95°C for 5 minutes in elution buffer containing 1% SDS and 20 mM TRIS (pH 7.5). Eluted proteins were collected from the supernatant and stored at –80°C until analysis by liquid chromatography – mass spectrometry, as detailed in the Supplemental Methods.

### Use of generative artificial intelligence

During the preparation of this manuscript, the authors used ChatGPT (OpenAI), based on the GPT-5.2 large language model (February 2026 version), for language editing. The tool was used solely to refine wording and grammar. All scientific content was developed, reviewed, and approved by the authors, who take full responsibility for the manuscript.

### Data, materials, and software availability

Mass spectrometry data have been deposited to the ProteomeXchange Consortium (PXD074429) via the jPOST partner repository (JPST004302) (*88*). All other data supporting the findings of this study are included in the article and its supplemental materials. DNA constructs and transgenic parasite lines used in this study are available upon reasonable request.

## ACKNOWLEDGEMENTS

This work was funded by the Bundesministerium für Forschung, Technologie und Raumfahrt (BMFTR; Federal Ministry of Research, Technology and Space; grant number 01KI2104 to JMM). Responsibility for the content of this publication lies solely with the authors. We also acknowledge financial support from the priority program SPP 2225 “Exit Strategies of Intracellular Pathogens” of the Deutsche Forschungsgemeinschaft (DFG; German Research Foundation; projects 531646514 to JMM and 446605368 to UD). We thank Tobias Spielmann from the Bernhard Nocht Institute for Tropical Medicine in Hamburg and Dave Richards from the Université Laval in Québec for permission to use the P40PX-mSc, GBP130_1-180_-mSc, mSc-Rab5a, and mSc-Rab7 plasmids.

## AUTHOR CONTRIBUTIONS

Conceptualization: JMM

Methodology: YS, MS, CS, TZ, FH, UD, RR, JMM

Investigation: YS, MS, CS, TZ, FH, RR, JMM

Data Curation and Analysis: YS, TZ, UD, RR, JMM

Funding Acquisition: UD, JMM

Writing – Original Draft: YS, JMM

Writing – Review & Editing: YS, MS, CS, TZ, FH, UD, RR, JMM

## Supplemental Material

## SUPPLEMENTAL METHODS

### Proteolytic digestion of immunoprecipitated proteins

Eluted proteins were digested on a Biomek i7 robotic pipetting platform (Beckman Coulter Life Sciences) using an adapted single-pot solid-phase-enhanced sample preparation protocol (*89*). Digestion steps followed the protocol from Distler et al., 2021 (*90*) with minor modifications. Proteins were reduced with 20 mM dithiothreitol (DTT) for 30 minutes at 45°C, then alkylated with iodoacetamide (IAA) for 30 minutes at room temperature, and excess IAA was quenched with DTT. Afterwards, 2.5 µg of carboxylate-modified paramagnetic beads (GE Healthcare) were added, as described previously (*89*). Samples were adjusted to a final concentration of 70% acetonitrile and incubated with gentle shaking for 20 minutes. Beads were immobilized by incubation on a magnetic rack for 2 minutes, washed four times with 70% ethanol and once with acetonitrile (ACN), and resuspended in 50 mM NH_4_HCO_3_ containing 0.4 µg trypsin (Promega). Digestion proceeded overnight at 37°C and was stopped by acidification with trifluoroacetic acid (TFA).

Peptides were desalted using 25 mg Sep-Pak tC18 µElution plates (Waters). Solid-phase extraction resin was conditioned with 200 µL methanol, followed by rinses with 200 µL of 80% ACN and 0.1% TFA. Peptides were then loaded onto the C18 columns followed by washing with 1 mL of 0.1% TFA. Peptides were eluted with 50 µL of 80% ACN and 0.1% TFA. Desalted peptides were frozen at –80°C, lyophilized, and subsequently resuspended in 20 µL 0.1% formic acid (FA) for LC-MS analysis.

### Liquid chromatography – mass spectrometry

Samples were analyzed using a nanoElute liquid chromatography system coupled online to a timsTOF Pro mass spectrometer (both from Bruker). Peptides were separated at 400 nL per minute on a heated (50°C) reversed-phase Aurora Ultimate C18 UHPLC emitter column (25 cm x 75 µm, 1.7 µm, IonOpticks) in direct injection mode at 600 bar. Mobile phase A was 0.1% FA in water, and mobile phase B 0.1% FA in ACN. Separation was achieved with a linear gradient from 2 to 37% B over 39 minutes, followed by a 5-minute rinse at 95% B. Eluting peptides were analyzed in positive mode ESI-MS using parallel accumulation serial fragmentation enhanced data-independent acquisition (diaPASEF) (*91*). The dual TIMS operated at near 100% duty cycle with equal accumulation and ramp times of 100 ms each across a mobility range of 1/K_0_ = 0.6-1.6 Vs cm^-2^. A total of 36 x 25 Th isolation windows from *m/z* 300-1,165 were defined, yielding 2-3 diaPASEF scans per TIMS cycle and an overall cycle time of 1.7 seconds. Collision energy was ramped linearly with mobility from 59 eV at 1/K_0_ = 1.3 Vs cm^-2^ to 20 eV at 1/K_0_ = 0.85 Vs cm^-2^. Samples were analyzed in triplicate.

### Processing of raw mass spectrometry data

Raw data were processed using DIA-NN (version 2.2.0) (*92*) with default settings for library-free database search. A custom database was compiled including the UniProtKB *P. falciparum* reference proteome (March 2025 release, 5,495 entries) and common contaminants. Trypsin was specified as the protease, allowing one missed cleavage. Carbamidomethylation was set as a fixed modification, with no variable modifications allowed. Peptide lengths were restricted to 7-30 amino acids. The precursor and product ion *m/z* ranges were 300-1,800 and 200-1,800, respectively. Quantification was performed in “QuantUMS (high precision)” mode without normalization. MS1 and MS2 mass accuracies, as well as scan window sizes, were optimized using DIA-NN’s built-in algorithms, maintaining peptide precursor false discovery rates below 1%.

### Statistical downstream analysis of proteomic data

Proteins had to be identified by at least two peptides to be included in the final dataset (Supplemental Dataset 1). Statistical analysis was performed using Student’s t-test, with multiple hypothesis testing corrected by the Benjamini-Hochberg method at a false discovery rate of 1%. Identified proteins were ranked based on a combined score reflecting both enrichment and statistical significance, calculated as log_2_ fold-change multiplied by the negative log_10_ of the p-value. To account for the higher protein content in the *Pf*ATG18 group relative to wild-type controls, proteins detected exclusively in the *Pf*ATG18 pull-downs were excluded from ranking.

**Supplemental Figure S1.**
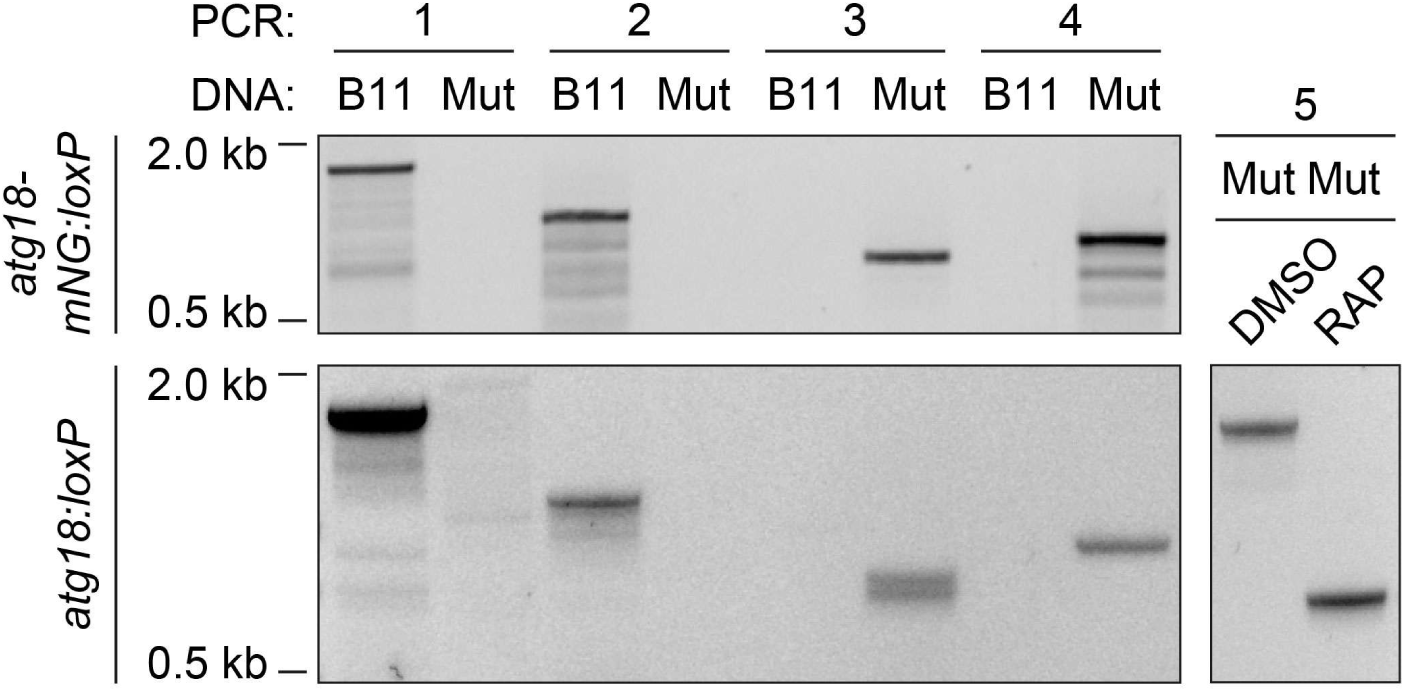
Validation of *Pf*ATG18 DiCre mutants by diagnostic PCR. Diagnostic PCRs, as indicated in Figure 1B, were performed using the primer combinations listed in Supplemental Table S1. Genomic DNA from parental B11 parasites, *atg18-mNG:loxP* and *atg18:loxP* mutants (Mut) was used as template. The excision-specific PCR5 used to analyze *atg18-mNG:loxP* parasites is shown in Figure 2A.

**Supplemental Figure S2.**
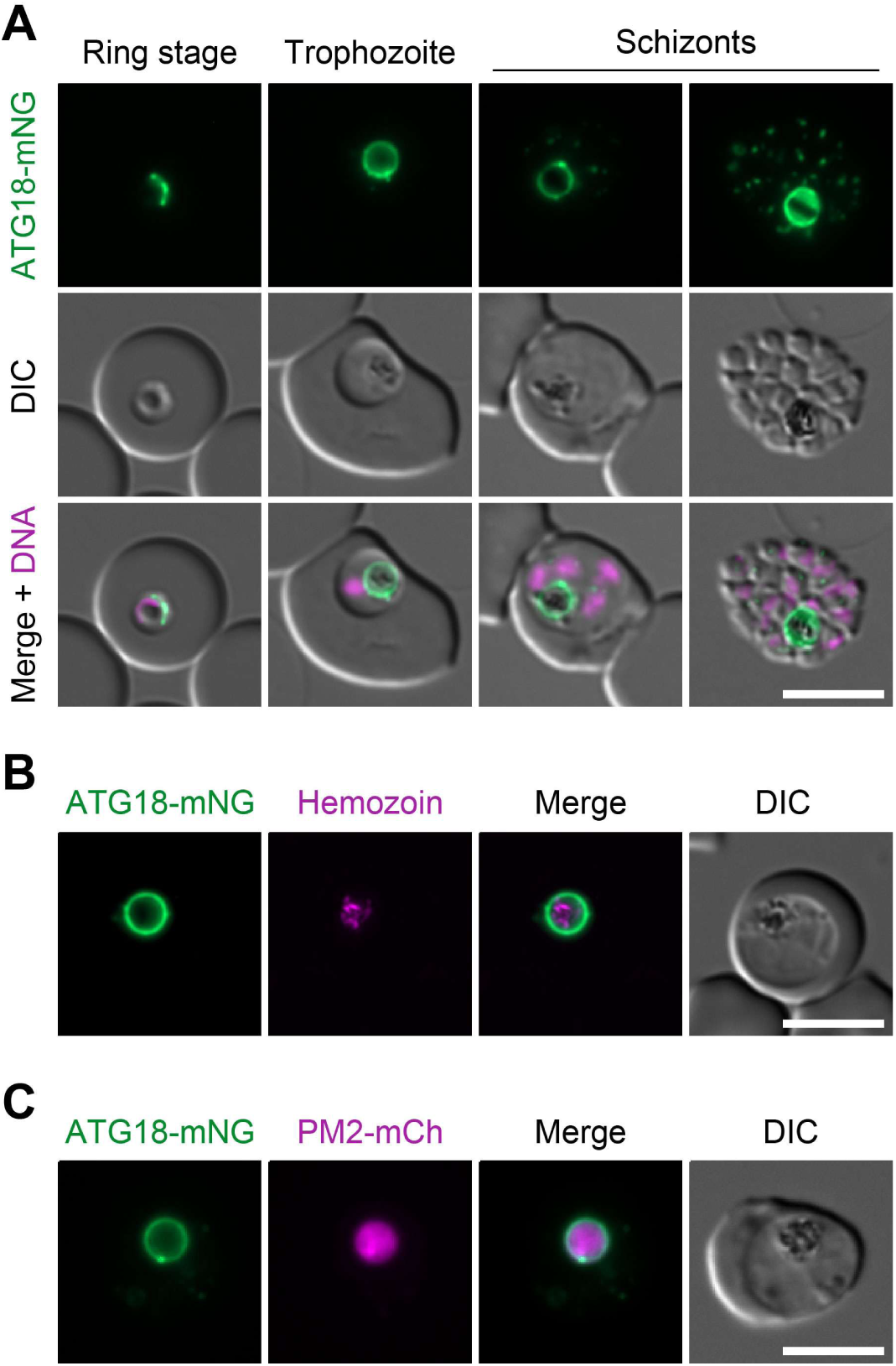
Expression and localization of *Pf*ATG18 as observed by live fluorescence microscopy. **(A)** *Pf*ATG18 is expressed throughout the entire asexual intraerythrocytic cycle. Representative parasite stages are shown. DNA: Hoechst 33342. **(B)** *Pf*ATG18 localizes around hemozoin, visualized using two crossed polarizers. **(C)** *Pf*ATG18-mNG staining delineates the DV, as demonstrated by co-localization with PM2-mCh. Scale bars: 5 µm.

**Supplemental Figure S3.**
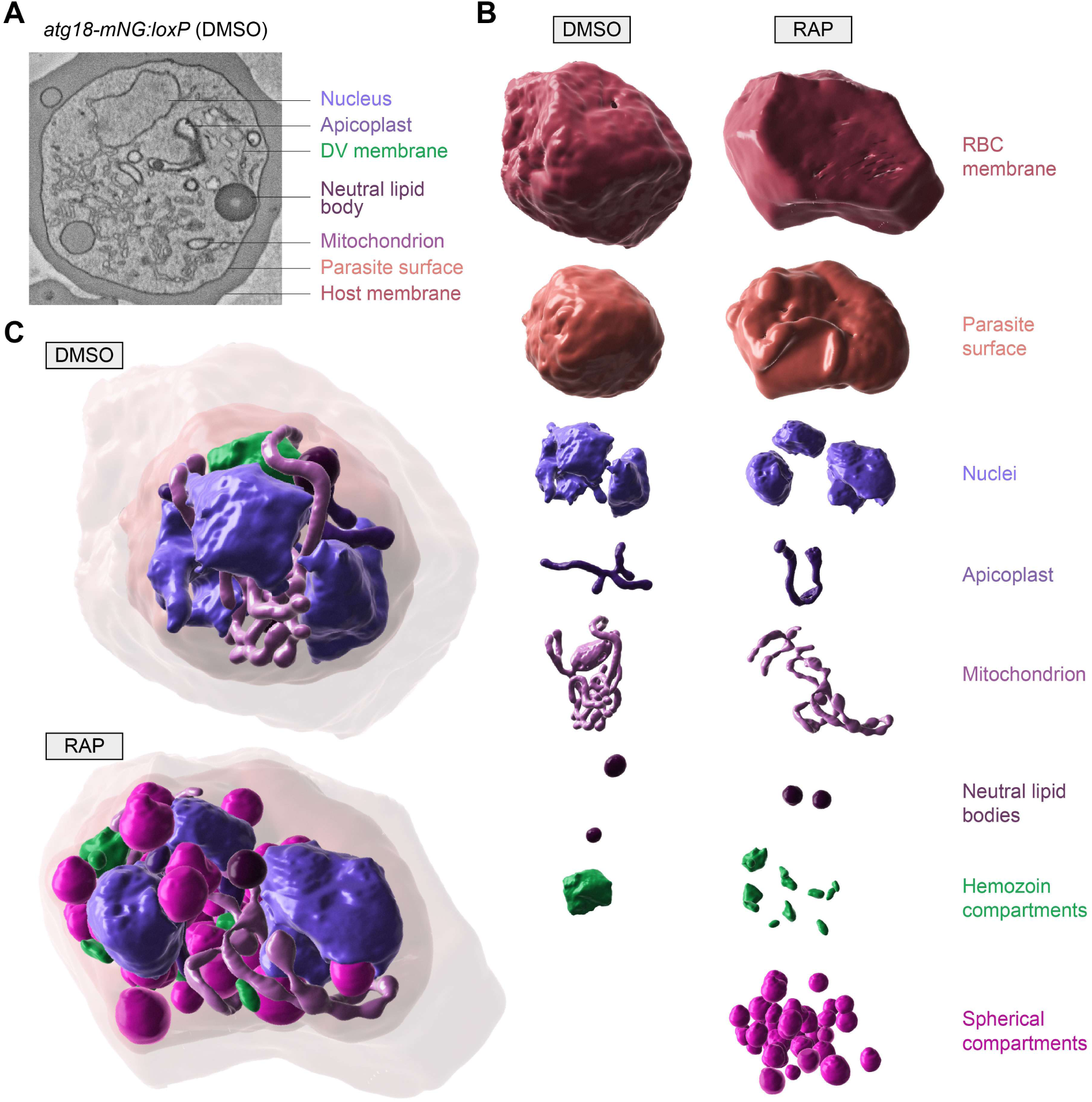
SBF-SEM analysis of *atg18-mNG:loxP* parasite architecture. *Pf*ATG18 loss was induced at the ring stage and parasites were analyzed at 36 hpi. **(A)** Representative SBF-SEM section of a DMSO control parasite, highlighting subcellular features selected for segmentation. The apicoplast was readily distinguished from the mitochondrion by its stronger membrane contrast. **(B, C)** 3D reconstructions of parasite structures in DMSO– and RAP-treated cells, shown individually (B) or combined (C). See also Supplemental Movies S1-S4.

**Supplemental Figure S4.**
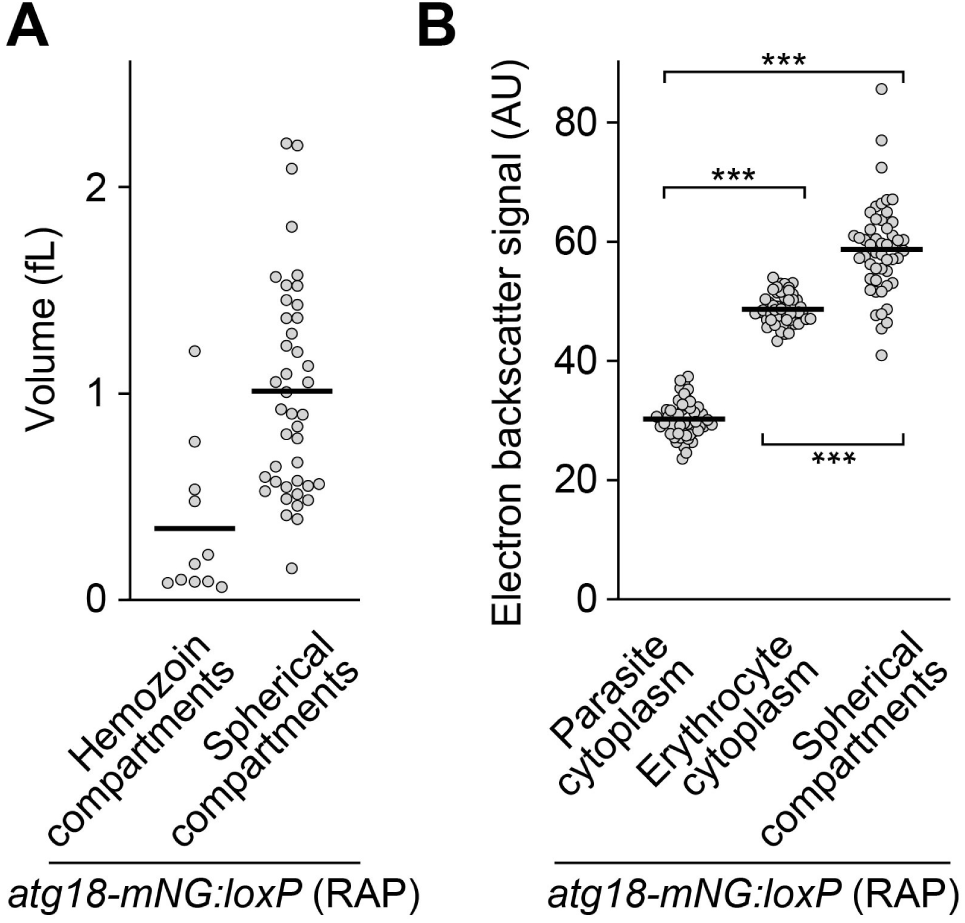
Volume and electron contrast of subcellular compartments in *Pf*ATG18-deficient parasites analyzed by SBF-SEM. **(A)** Volumes of hemozoin-containing DV fragments and spherical compartments. n = 11 DV fragments and 42 spherical compartments from 1 reconstructed parasite. **(B)** Electron backscatter signals of parasite cytoplasm, erythrocyte cytoplasm and spherical compartments. Individual and mean values; one-way ANOVA with Tukey’s test; n = 50 recordings from 5 infected cells. ***, P < 0.001.

**Supplemental Figure S5.**
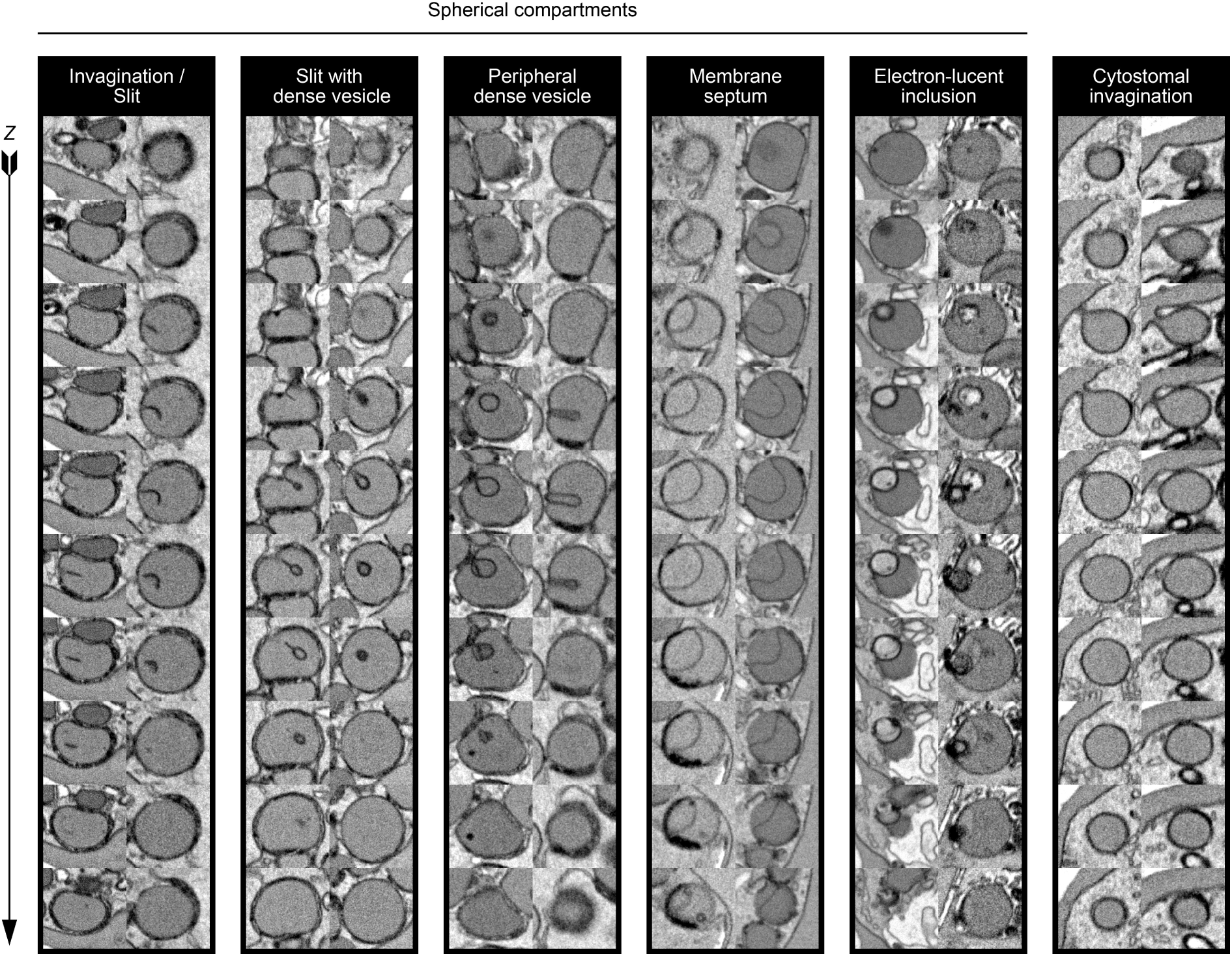
Spherical compartments in *Pf*ATG18-deficient parasites exhibit diverse membrane features. Representative SBF-SEM Z-stacks of individual spherical compartments and cytostomes are shown, with two examples for each morphology. Note that electron-lucent inclusions were rare and that cytostomes lacked internal membrane profiles.

**Supplemental Figure S6.**
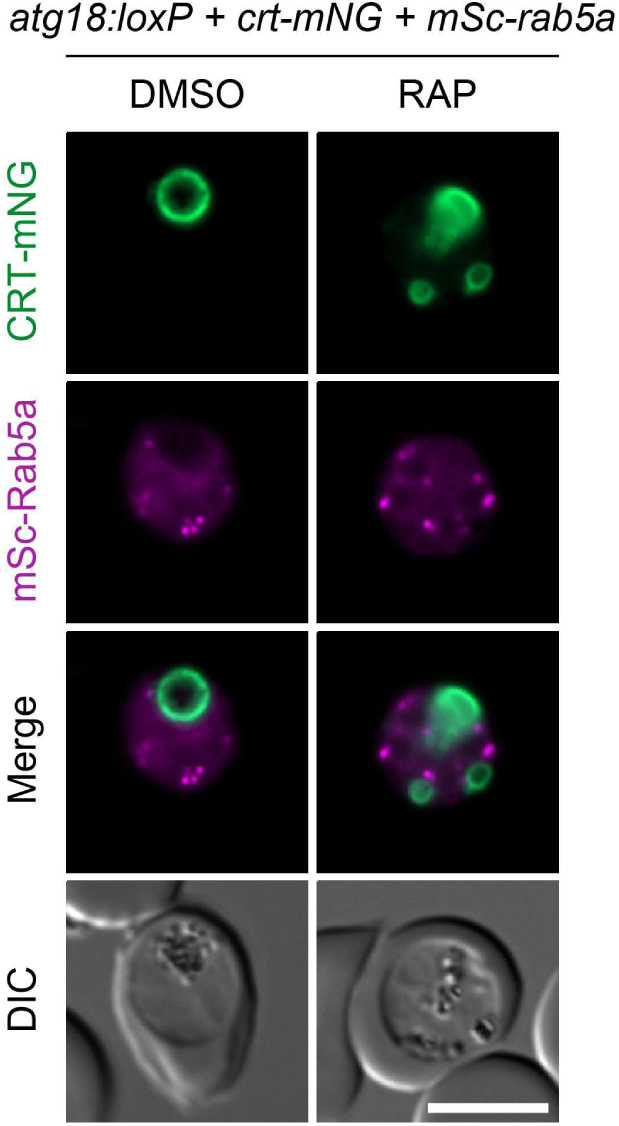
Unaltered Rab5A localization in *Pf*ATG18-deficient parasites. Live fluorescence micrographs of *atg18:loxP* parasites expressing endogenous CRT-mNG and episomal mSc-Rab5A. Scale bar: 5 µm.

**Supplemental Figure S7.**
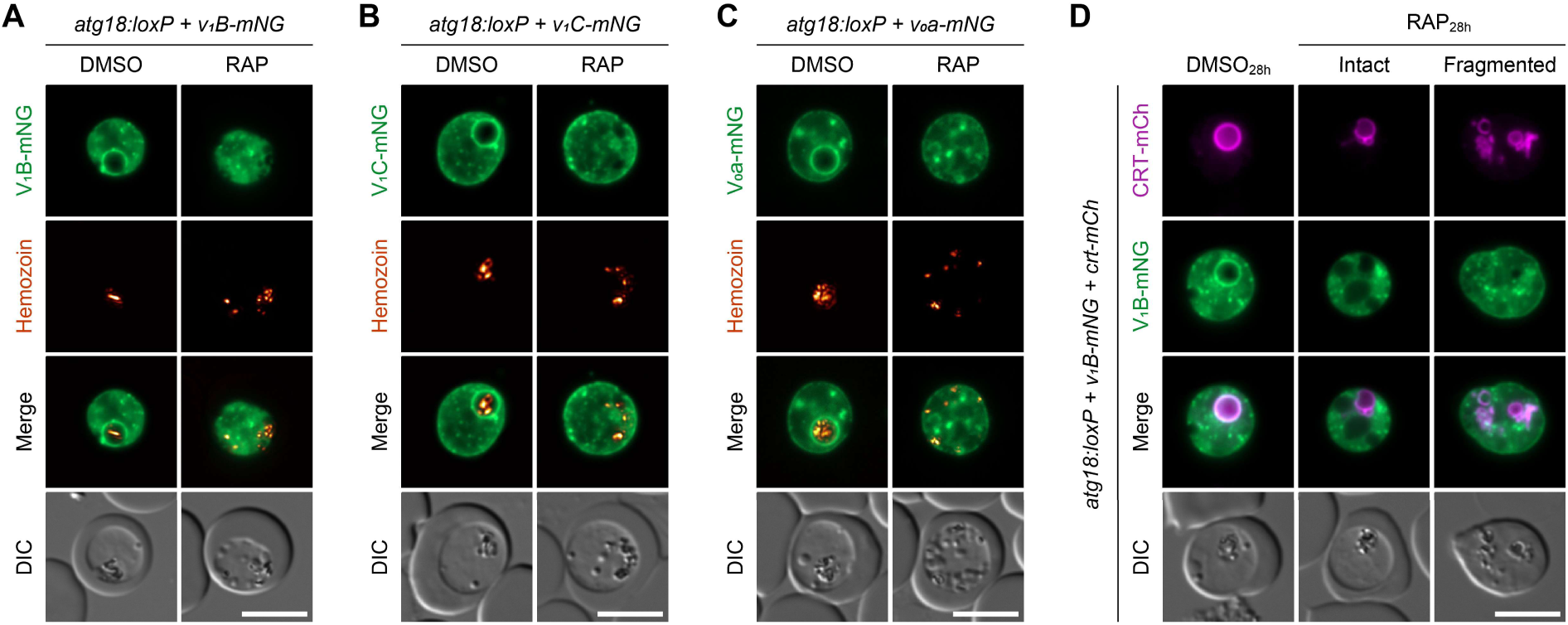
V-ATPase integrity is compromised upon loss of *Pf*ATG18, preceding DV fission. **(A-C)** Live-cell imaging of mNG-tagged V-ATPase subunits V_1_B (A), V_1_C (B), and V_0_a (C) reveals widespread V-ATPase disengagement from the DVM. Hemozoin was visualized using two crossed polarizers. **(D)** Colocalization analysis of CRT-mCh and V_1_B-mNG at the onset of vacuolar fragmentation. *Pf*ATG18 knockout was induced at 28 hpi, and parasites were analyzed 8 hours later. Note that V_1_B is already dissociated in *Pf*ATG18-deficient parasites, regardless of whether the DV is intact or already fragmented. Scale bars: 5 µm.

**Supplemental Table S1.**
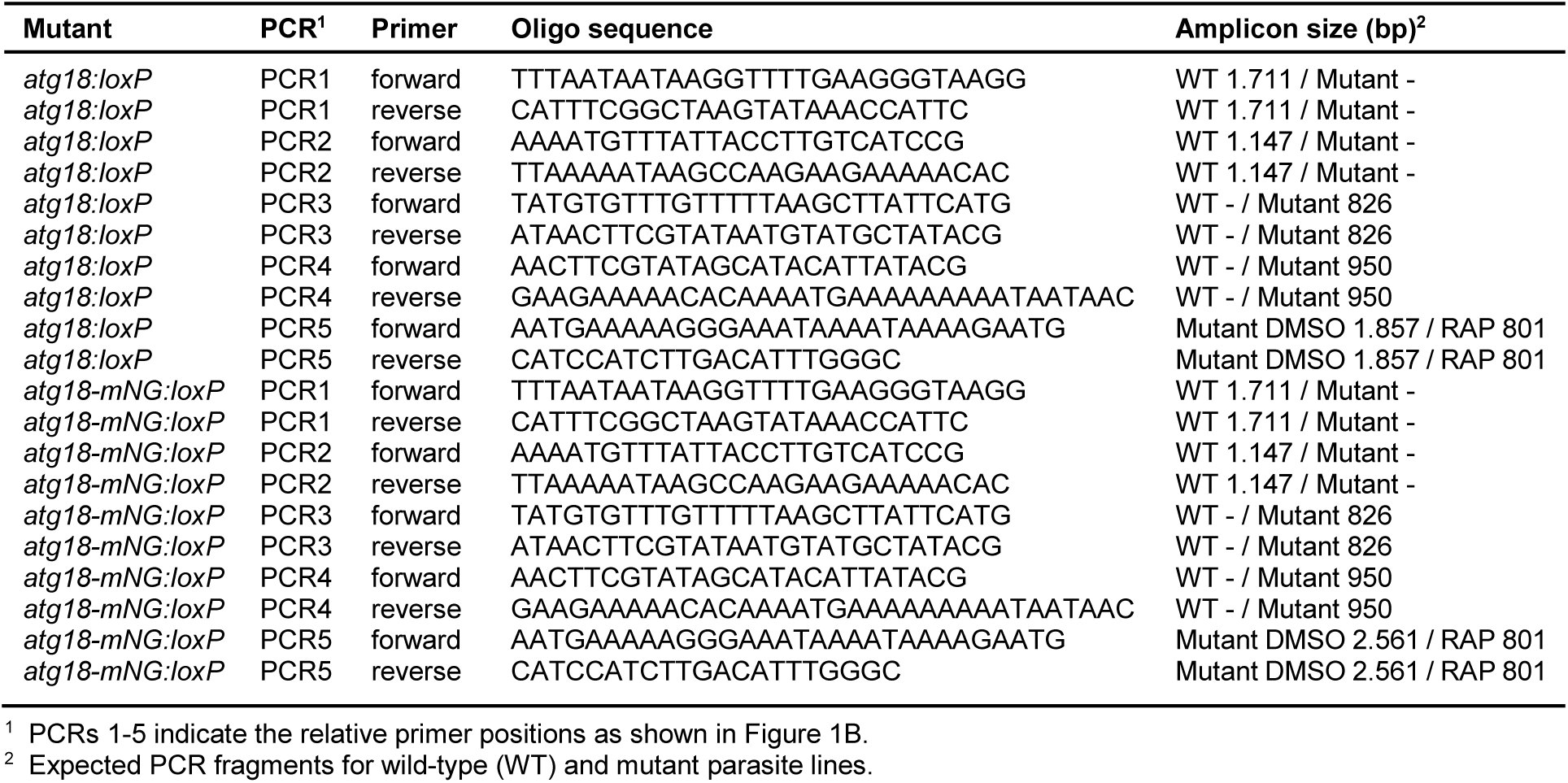
Diagnostic PCRs performed in this study.

**Supplemental Table S2.**
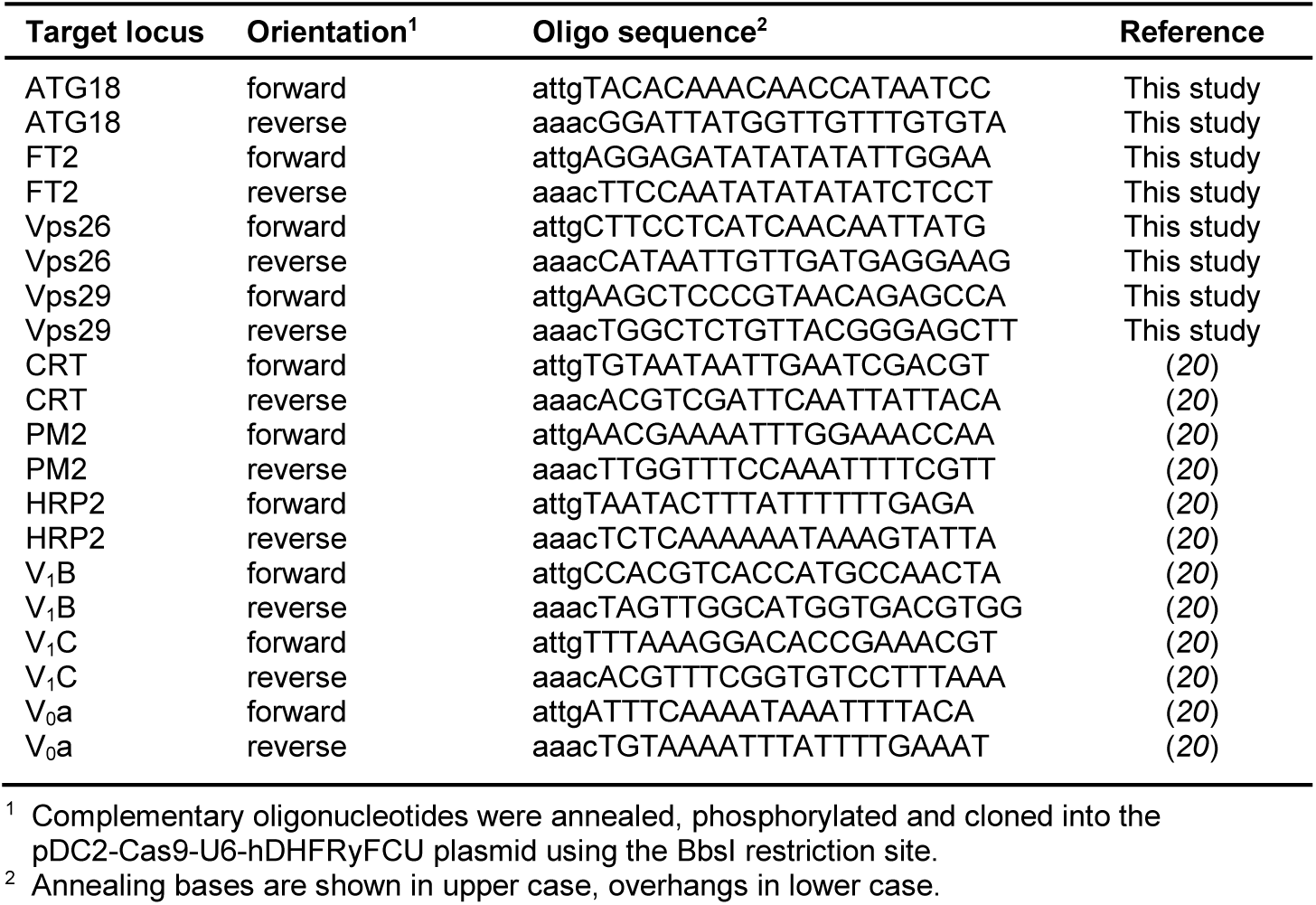
Oligonucleotides used for guide plasmid generation.

**Supplemental Table S3.**
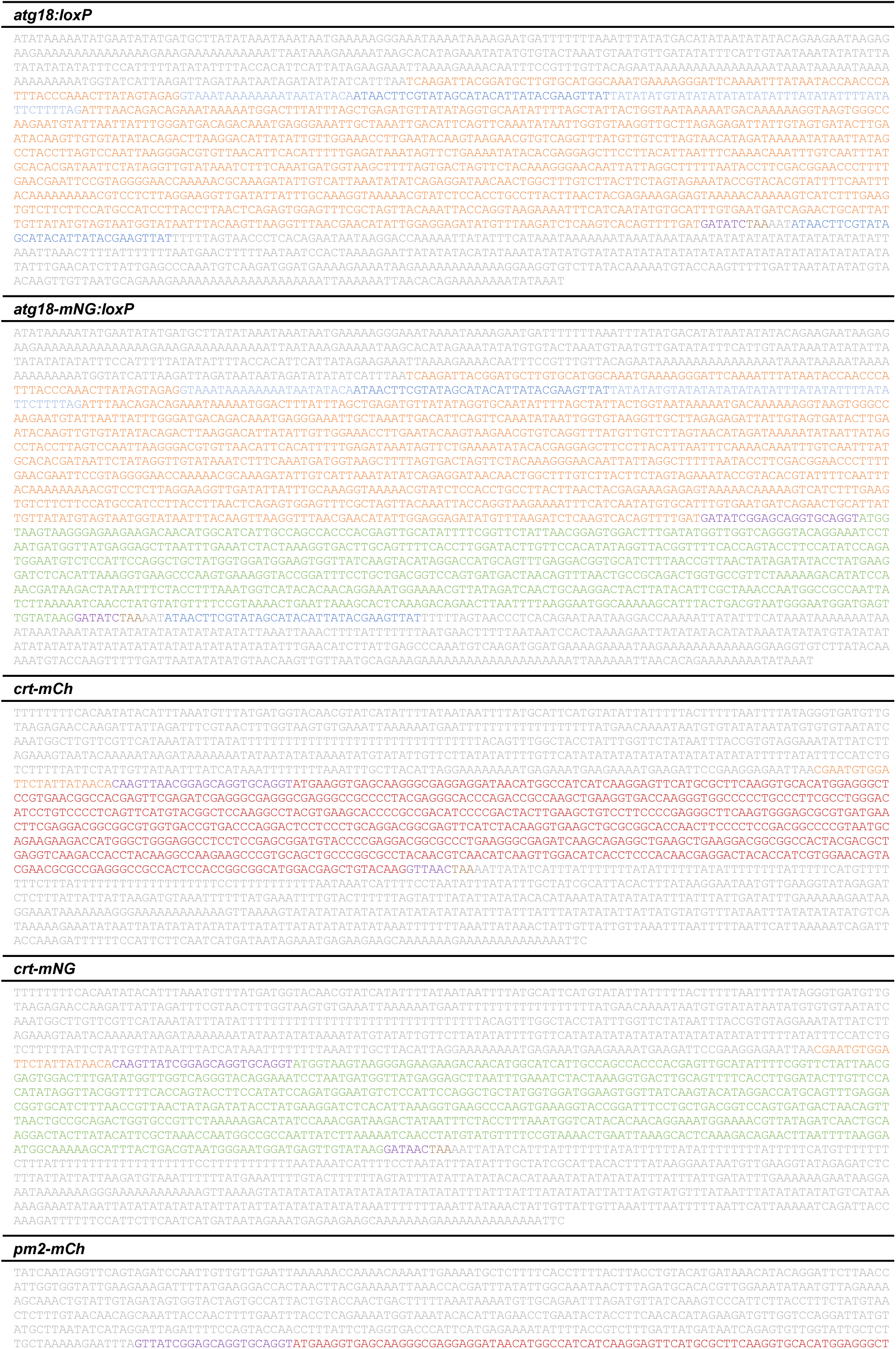
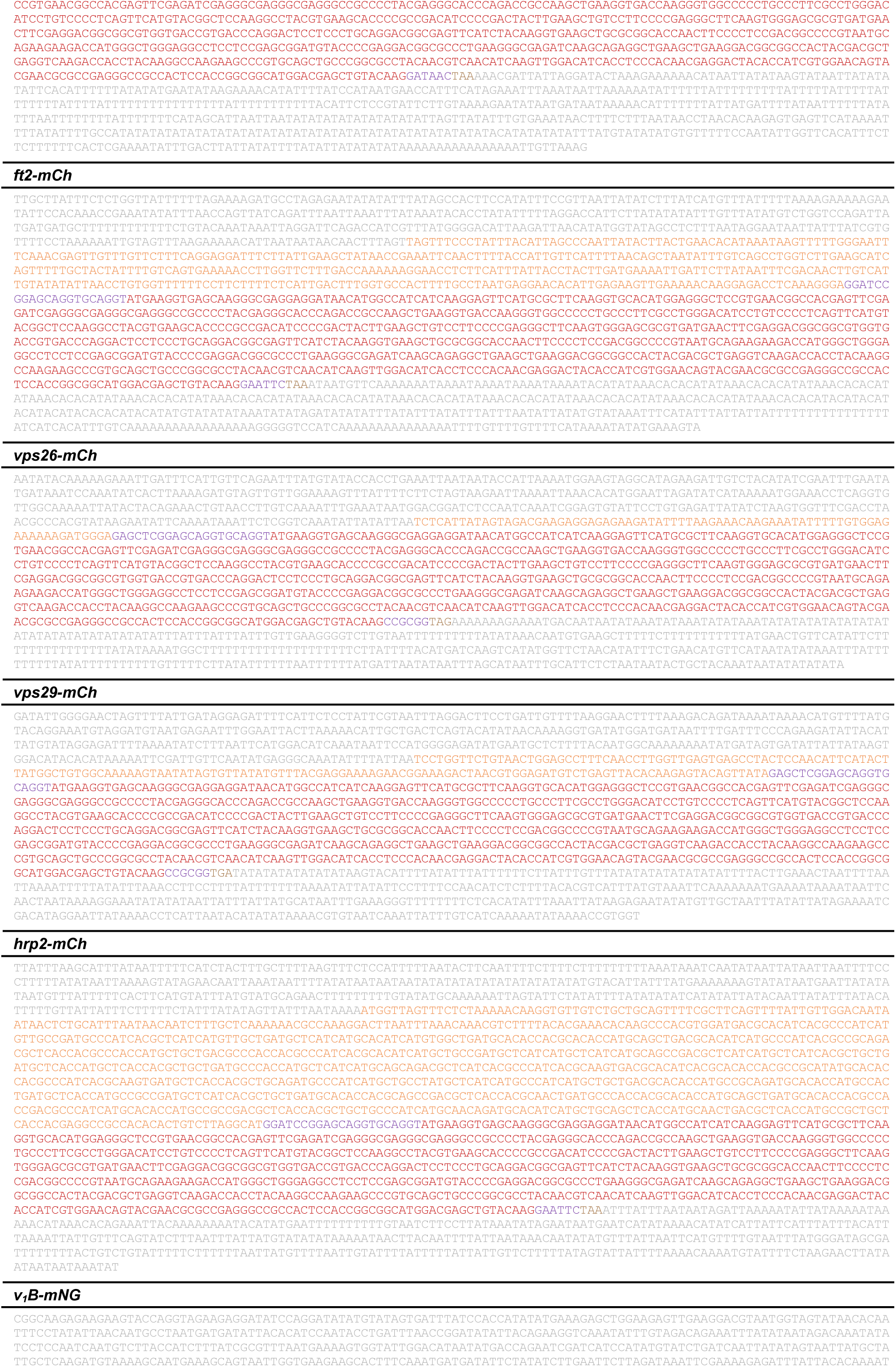
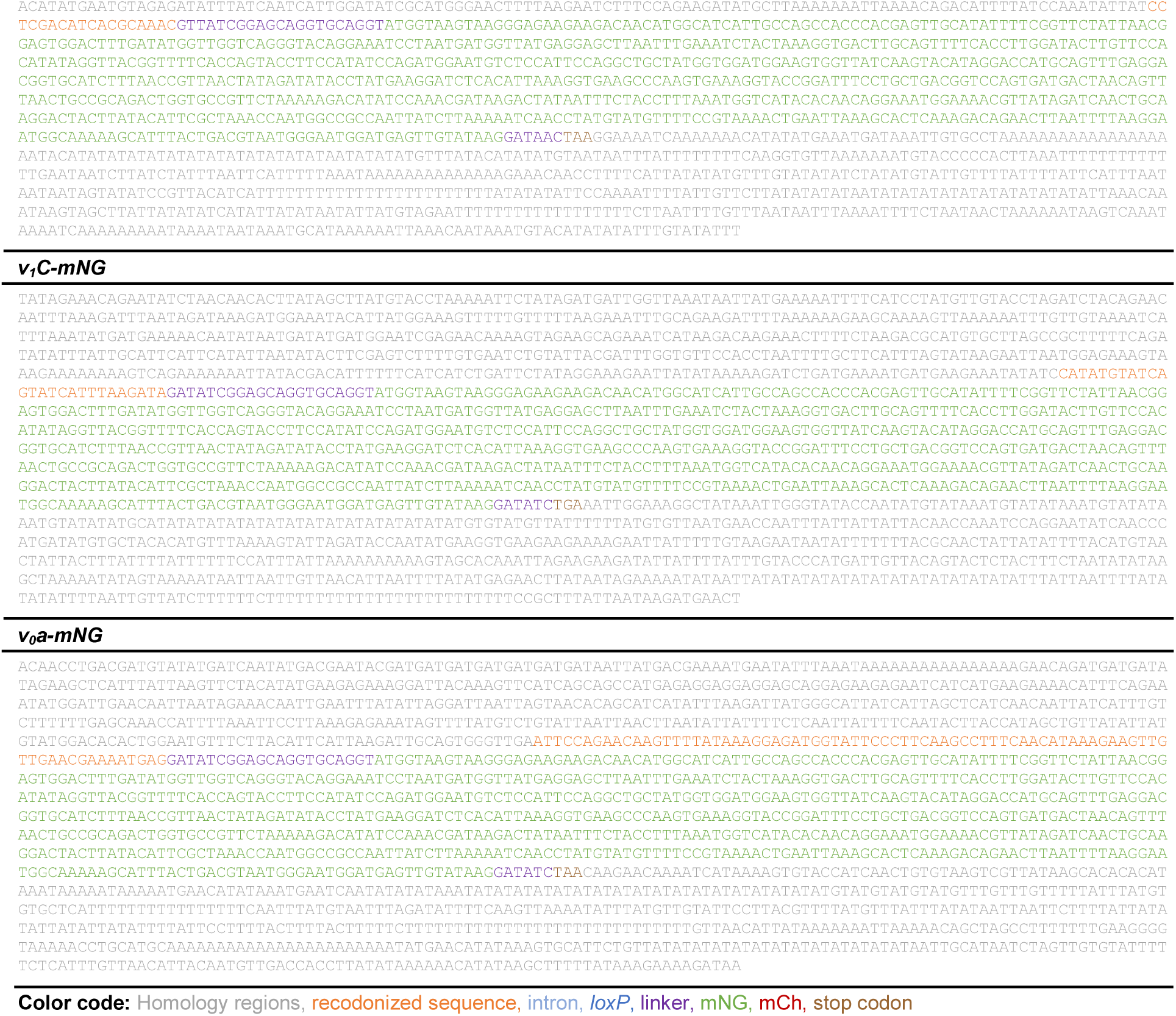
Repair templates used in this study.

**Supplemental Table S4.** Transgenic parasite lines used in this study.

| Mutant | Parental strain<sup>1</sup> | Description |
| --- | --- | --- |
| B11 | 3D7 | Constitutive expression of DiCre recombinase |
| *atg18:loxP* | B11 (47) | Conditional deletion of untagged ATG18 by DiCre |
| *atg18-mNG:loxP* | B11 (47) | Conditional deletion of mNG-tagged ATG18 by DiCre |
| *atg18:loxP + crt-mNG* | *atg18:loxP* | Endogenous mNG-tagging of CRT in ATG18 DiCre line |
| *atg18:loxP + v<sub>1</sub>B-mNG* | *atg18:loxP* | Endogenous mNG-tagging of V<sub>1</sub>B in ATG18 DiCre line |
| *atg18:loxP + v<sub>1</sub>C-mNG* | *atg18:loxP* | Endogenous mNG-tagging of V<sub>1</sub>C in ATG18 DiCre line |
| *atg18:loxP + v<sub>0</sub>a-mNG* | *atg18:loxP* | Endogenous mNG-tagging of V<sub>0</sub>a in ATG18 DiCre line |
| *atg18:loxP + crt-mNG + p40px-mSc* | *atg18:loxP + crt-mNG* | Episomal expression of mSc-tagged P40PX in ATG18 DiCre line with mNG-tagged CRT |
| *atg18:loxP + crt-mNG + mSc-rab7* | *atg18:loxP + crt-mNG* | Episomal expression of mSc-tagged Rab7 in ATG18 DiCre line with mNG-tagged CRT |
| *atg18:loxP + v<sub>1</sub>B-mNG + crt-mCh* | *atg18:loxP + v<sub>1</sub>B-mNG* | Endogenous mCh-tagging of CRT in ATG18 DiCre line with mNG-tagged V<sub>1</sub>B |
| *atg18:loxP + v<sub>0</sub>a-mNG + crt-mCh* | *atg18:loxP + v<sub>0</sub>a-mNG* | Endogenous mCh-tagging of CRT in ATG18 DiCre line with mNG-tagged V<sub>0</sub>a |
| *atg18-mNG:loxP + crt-mCh* | *atg18-mNG:loxP* | Endogenous mCh-tagging of CRT in ATG18-mNG DiCre line |
| *atg18-mNG:loxP + vps26-mCh* | *atg18-mNG:loxP* | Endogenous mCh-tagging of Vps26 in ATG18-mNG DiCre line |
| *atg18-mNG:loxP + vps29-mCh* | *atg18-mNG:loxP* | Endogenous mCh-tagging of Vps29 in ATG18-mNG DiCre line |
| *atg18-mNG:loxP + ft2-mCh* | *atg18-mNG:loxP* | Endogenous mCh-tagging of FT2 in ATG18-mNG DiCre line |
| *atg18-mNG:loxP + pm2-mCh* | *atg18-mNG:loxP* | Endogenous mCh-tagging of PM2 in ATG18-mNG DiCre line |
| *atg18-mNG:loxP + hrp2-mCh* | *atg18-mNG:loxP* | Endogenous mCh-tagging of HRP2 in ATG18-mNG DiCre line |
| *atg18-mNG:loxP + mSc-rab5a* | *atg18-mNG:loxP* | Episomal expression of mSc-tagged Rab5a in ATG18-mNG DiCre line |
| *atg18-mNG:loxP + mSc-rab7* | *atg18-mNG:loxP* | Episomal expression of mSc-tagged Rab7 in ATG18-mNG DiCre line |
| *v<sub>0</sub>c:loxP + crt-mNG + gbp130<sub>1-180</sub>-mSc* | *v<sub>0</sub>c:loxP + crt-mNG (20)* | Episomal expression of mSc-tagged GBP130<sub>1-180</sub> in V<sub>0</sub>c DiCre line with CRT-mNG |

<sup>1</sup> Recipient strains used as genetic background for transfections.

**Supplemental Movies S1-S4. Ultrastructural analysis of *atg18-mNG:loxP* parasites by SBF-SEM**

*Pf*ATG18 deletion was induced at the ring stage, and parasites were analyzed at 36 hpi.

**(S1)** Z-stack of a DMSO-treated control parasite.

**(S2)** Z-stack of a RAP-treated parasite.

**(S3)** 3D reconstruction of a DMSO-treated control parasite.

**(S4)** 3D reconstruction of a RAP-treated parasite.

Color coding of individual structures corresponds to that in Supplemental Figure S3.

